# Genetic screens reveal a central role for heme biosynthesis in artemisinin susceptibility

**DOI:** 10.1101/746974

**Authors:** Clare R. Harding, Saima M. Sidik, Boryana Petrova, Nina F. Gnädig, John Okombo, Kurt E. Ward, Benedikt M. Markus, David A. Fidock, Sebastian Lourido

## Abstract

Artemisinins have revolutionized the treatment of *Plasmodium falciparum* malaria, however, resistance threatens to undermine global control efforts. To explore artemisinin resistance in apicomplexan parasites broadly, we used genome-scale CRISPR screens recently developed for *Toxoplasma gondii* to discover sensitizing and desensitizing mutations. Using a sublethal concentration of dihydroartemisinin (DHA), we uncovered the putative porphyrin transporter Tmem14c whose disruption increases DHA susceptibility. Screens performed under high doses of DHA provided evidence that mitochondrial metabolism can modulate resistance. We show that disruption of a top candidate from the screens, the mitochondrial protease DegP2, lowered levels of free heme and decreased DHA susceptibility, without significantly altering fitness in culture. Deletion of the homologous gene in *P. falciparum, PfDegP*, similarly lowered heme levels and DHA susceptibility. These results expose the vulnerability of the heme biosynthetic pathway for genetic perturbations that can lead to survival in the presence of DHA. We go on to show that chemically reducing heme biosynthesis can decrease the sensitivity of both *T. gondii* and *P. falciparum* to DHA, suggesting guidelines for developing combination therapies.

## INTRODUCTION

Since their discovery and characterization as potent antimalarials, artemisinin (ART) and its synthetic derivatives have emerged as important drugs in treating infectious diseases and are being actively investigated in cancer therapy^1,2^. Notably, artemisinin combination therapies (ACTs) have transformed malaria treatment, particularly in response to widespread resistance to previous classes of anti-parasitic compounds. Artemisinins require activation within cells by scission of an endoperoxide bridge through the Fe^2+^ center of heme^1,3^. Cleavage of this bond results in the production of ART radicals, which react with hundreds of proteins^4,5^, lipids^6^, and metabolites,^7^ quickly leading to cell death.

Artemisinins are labile, and their short half lives (∼1 h in humans) lead modest, stage-specific reductions in parasite sensitivity to result in treatment failure. Such treatment failure was first noted in Southeast Asia a decade ago, and is increasingly frequent^8–11^. Treatment failure has been associated with genetic polymorphisms in *P. falciparum* Kelch13 (K13), most notably K13^C580Y 12,13^. The mechanism underlying this effect remains under scrutiny, but it is thought to involve changes in the unfolded protein response, autophagy, and PI3K activity^14–17^. Decreased ART sensitivity has also been attributed to mutations unrelated to K13, such as coronin^18^, clathrin adaptors^19–21^, and cysteine proteases^22^. *P. falciparum* strains without identified causative mutations have displayed decreased clearance as well^23,24^. Hence, a unified picture of ART sensitivity remains to be elucidated.

Directed evolution coupled with whole-genome sequencing has identified targets and resistance pathways for many antiparasitic compounds^18,25–27^. Such approaches can be time-consuming, may fail to detect minor mutations or those that negatively affect parasite fitness, and can only be used for positive selection schemes. Recently, whole-genome CRISPR-based mutational approaches have uncovered drug-resistance mechanisms in cancer^28–30^. We have shown that CRISPR screens in *T. gondii* provide a comparatively fast method to identify causal mutations using both positive and negative selection^31,32^.

Here, we establish that analogous mutations in *T. gondii* and *P. falciparum* can affect sensitivity to the active ART metabolite, DHA. We demonstrate that a mutation homologous to the canonical *P. falciparum* K13^C580Y^ similarly reduced *T. gondii* sensitivity to DHA. Genome-wide screens in *T. gondii* further identified mutants in a porphyrin transporter (Tmem14c) that are more susceptible to DHA, as well as mutations in several genes involved in mitochondrial metabolism that decreased drug sensitivity. Our results point to a central role for heme availability in limiting DHA activation and determining *T. gondii* sensitivity to the compound. Critically, a mitochondrial protease (DegP2) whose disruption decreases free-heme levels, can be mutated to decrease DHA sensitivity without impacting parasite fitness. We extended these results to *P. falciparum*, where we showed that impairing heme biosynthesis, chemically or by deleting the *P. falciparum* homolog of *DegP2*, also decreases DHA sensitivity. Taken together with our *T. gondii* data, these results suggest that minimal genetic perturbations can alter levels of free heme in both genera enhancing the survival of mutants in the presence of DHA.

## RESULTS

### Using *T. gondii* to understand apicomplexan DHA sensitivity

To establish the timing and concentrations under which to assess changes in *T. gondii* sensitivity to DHA, we treated intracellular parasites with varying concentrations of DHA or pyrimethamine—the front-line treatment for *T. gondii*—for 24 hours and measured parasite viability by counting the number of vacuoles containing two or more parasites in vehicle or drug treated conditions. Both drugs completely blocked parasite replication at the highest doses (**Figure 1a**), recapitulating the previously reported IC_50_ for pyrimethamine^33^ and underscoring the potency of DHA against *T. gondii.*

**Figure 1.**
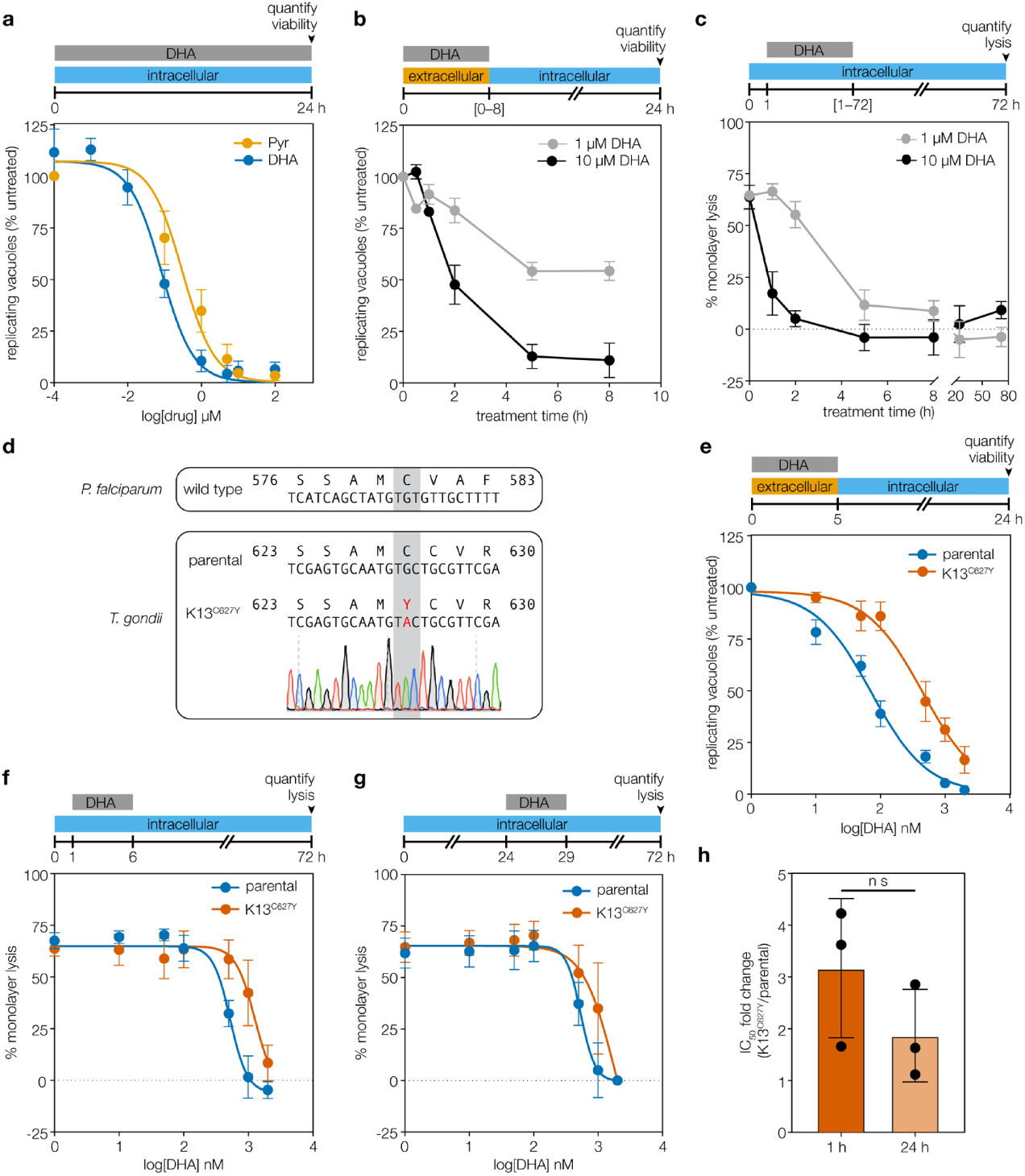
DHA kills wild-type *T. gondii* efficiently and its effect is influenced by mutation of K13. **a**, Treatment with increasing concentrations of dihydroartemisinin (DHA) or pyrimethamine (Pyr) over 24 h resulted in a reduction in parasite viability, measured as the number of vacuoles with two or more parasites normalized to untreated wells. Results are mean ± SEM for n = 4 independent experiments. **b**, Viability was similarly assessed for extracellular parasites treated with 1 or 10 µM DHA for varying periods of time, then washed and allowed to invade host cells. Viable vacuoles were normalized to untreated parasites kept extracellular for equal periods of time. Results are mean ± SEM for n = 3 independent experiments, each performed in technical triplicate. **c**, Quantification of host monolayer lysis after treatment with 1 or 10 μM DHA for the indicated time, normalised to uninfected monolayers. Results are mean ± SEM for n = 4 independent experiments. **d**, Alignment of the mutated region of K13 in *P. falciparum* and *T. gondii* displaying the chromatogram for the K13^C627Y^ *T. gondii* line. **e**, Extracellular dose-response curve for parental or K13^C627Y^ parasites treated with DHA for 5 h. Results are mean ± SEM for n = 7 or 5 independent experiments with parental or K13^C627Y^ parasites, respectively. **f**, Monolayer lysis following infection with parental or K13^C627Y^ parasites for 1 h, prior to a 5 h treatment with varying concentrations of DHA. Results are mean ± SEM for n = 4 independent experiments. **g**, Monolayer lysis following infection with parental or K13^C627Y^ parasites for 24 h, prior to 5 h treatment with varying concentrations of DHA. Results are mean ± SEM for n = 4 or 3 independent experiments with parental or K13^C627Y^ parasites, respectively. **h**, Graph summarising the fold change in DHA IC_50_ between parental and K13^C627Y^ parasites, treated 1 h or 24 h after invasion. Results are mean ± SD for n = 3 independent experiments, *p* value calculated from unpaired *t* test.

To isolate DHA’s effect on parasites from effects on host cells, we determined the minimum treatment time required for DHA to reach maximum lethality using *T. gondii’s* extracellular stages. Parasites were treated for increasing lengths of time, washed, applied to host cells, and left to replicate for 24 h. DHA treatment at 1 μM and 10 μM blocked parasite viability maximally after 5 h (**Figure 1b**). We similarly determined the treatment window for intracellular parasites by adding DHA to cultures at 1 h post invasion (1 h p.i.), washing out the compound after different lengths of time, and assessing viability as monolayer lysis 72 h post invasion (**Figure 1c**). Based on these results, we used 5 h treatments for both intracellular and extracellular parasites^34^.

### Homologous Kelch13 mutations decrease DHA sensitivity in *P. falciparum* and *T. gondii*

In *P. falciparum*, point mutations in *Kelch13* (*K13*), like C580Y and R538T, correlate with delayed clearance and increased survival of ring-stage parasites^12,13,35,36^. *K13* is conserved among apicomplexans; however, its role in DHA sensitivity has not been examined in *T. gondii*. We made a C627Y mutation in the *T. gondii* ortholog of *K13* (TGGT1_262150), corresponding to *P. falciparum* C580Y^13^ (**Figure 1d**). Extracellular K13^C627Y^ parasites showed decreased sensitivity to DHA compared to the parental strain (Figure 1e, IC_50_ values provided in **Supplementary Table 1**), indicating that the effect of K13 mutations on DHA sensitivity transcends genera. As in *P. falciparum*, K13^C627Y^ parasites were sensitive to prolonged DHA exposure and did not form plaques under continuous DHA treatment (**Supplementary Figure 1**).

DHA susceptibility changes over the *P. falciparum* erythrocytic cycle, with trophozoites displaying greater sensitivity than rings or schizonts^37,38^. To determine whether active replication affects DHA sensitivity in *T. gondii*, we treated intracellular parasites immediately after invasion (**Figure 1f, Supplementary Table 1**)—before most parasites have initiated replication—or 24 h after invasion when most parasites are replicating^39^ (**Figure 1g, Supplementary Table 1**). K13^C627Y^ parasites were similarly resistant to DHA at both time points (Figure 1h) and we conclude that replication does not affect the reduced DHA sensitivity of K13^C627Y^ *T. gondii.*

### A genome-wide screen identifies mutants that are hypersensitive to DHA

We used CRISPR to generate a pool of mutant parasites, with ten gRNAs targeting each gene^31,32,40^, then propagated this pool in the presence or absence of a sublethal dose (50 nM) of DHA for 7 days (three passages for the untreated population; **Figure 2a**). Guides against a single gene (TGGT1_228110) were substantially depleted under drug treatment compared to the untreated control (**Figure 2b** and **Supplementary Table 2**). TGGT1_228110 encodes a previously unstudied *T. gondii* gene with two transmembrane domains, predicted to be dispensable under standard growth conditions^32^. Structural prediction using HHPred^41^ indicated that TGGT1_228110 is homologous to TMEM14c (*p* = 5 x 10^-21^)— a recently identified porphyrin transporter found in mammals^42^. Based on this homology, we named TGGT1_228110 Tmem14c.

**Figure 2.**
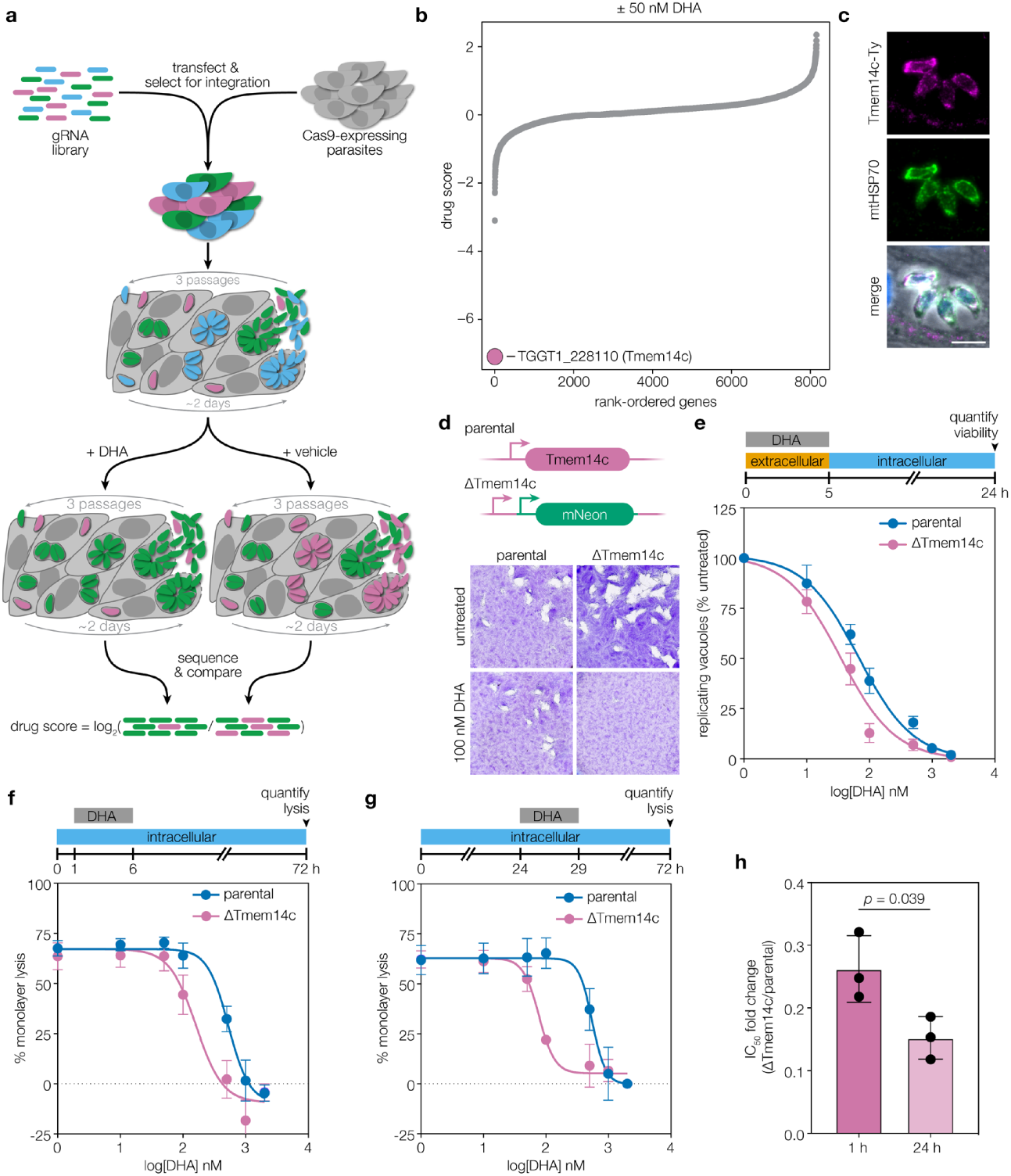
Genome-wide screen under sublethal DHA concentration reveals that loss of Tmem14c increases DHA sensitivity. **a**, Screening workflow. The drug score is defined as the log_2_-fold change in the relative gRNA abundance between the DHA-treated and vehicle populations, where lower scores are indicative of genes that enhance drug sensitivity when disrupted. **b**, Results of a genome-wide CRISPR screen comparing treatment with 50 nM DHA (equivalent to IC_5_) to vehicle-treated parasites. Guides against one gene, Tmem14c, were significantly depleted by the sublethal DHA concentration but retained in the vehicle control. **c**, Overexpression of Tmem14c-Ty co-localized with mitochondrial marker mtHSP70. Scale bar is 5 μm. **d**, ΔTmem14c parasites formed plaques normally under standard growth conditions, but their plaquing ability was attenuated in the presence of 100 nM DHA, whereas the parental strain grew normally in both conditions. **e**, Extracellular ΔTmem14c parasites had a decreased IC_50_ compared to the parental line (*p* = 0.0003, extra-sum-of-squares F test). Results are mean ± SEM for n = 7 independent experiments. **f**, Monolayer lysis following infection with parental or ΔTmem14c parasites for 1 h, prior to a 5 h treatment with varying concentrations of DHA. Results are mean ± SEM for n = 4 or 3 independent experiments for parental or ΔTmem14c parasites, respectively. **g**, Monolayer lysis following infection with parental or ΔTmem14c parasites for 24 h, prior to 5 h treatment with varying concentrations of DHA. Results are mean ± SEM for n = 4 independent experiments. **h**, Graph summarizing the fold change in DHA IC_50_ between the parental and ΔTmem14c parasites, treated 1 h or 24 h after invasion. Results are mean ± SD for 3 independent experiments, *p* value calculated from unpaired *t* test.

To examine the function of Tmem14c, we determined its subcellular localization and the effect of its deletion. A C-terminally Ty-tagged copy of Tmem14c localized to the parasite mitochondrion (**Figure 2c**). This agrees with the localization of the mammalian TMEM14C to the inner mitochondrial membrane^42^. We replaced the coding region of Tmem14c with an mNeonGreen expression cassette. (**Supplementary Figure 2**). Although ΔTmem14c grew normally under standard conditions, treating mutant parasites with 100 nM DHA for 7 days abolished plaque formation with minimal effects on the parental line (**Figure 2d**). We quantified the change in DHA sensitivity by treating extracellular parasites with varying concentrations of DHA, then quantifying their viability, and observed a modest increase in sensitivity for ΔTmem14c parasites compared to the parental line (**Figure 2e, Supplementary Table 1**). In contrast to the extracellular treatment used above, CRISPR screens maintain DHA pressure during parasite replication. We therefore assessed the DHA sensitivity of ΔTmem14c parasites during the intracellular cycle by treating with DHA for 5 h immediately after invasion (**Figure 2f, Supplementary Table 1**) or during replication (**Figure 2g, Supplementary Table 1**). In contrast to the parental and K13^C627Y^ parasite lines (**Figure 1h**), the sensitivity of ΔTmem14c to DHA was significantly more pronounced during replication (**Figure 2h**). This suggests that Tmem14c is more important in replicating parasites, which may require increased porphyrin biosynthesis.

### Genome-wide screens implicate TCA and heme biosynthesis pathways in DHA sensitivity

We performed two genome-wide CRISPR screens to identify mutations that decrease sensitivity to DHA. Following selection for integration of the gRNAs, parasite populations were treated with 0.5 μM DHA or vehicle, then cultured for the time required to passage the untreated population three times (approximately 6 days). The log_2_-fold change of gRNA abundance between the drug-treated and untreated populations (drug score) was used to rank all protein-coding genes (**Figure 3a, Supplementary Table 2**). Two replicates of this screen were pooled and analyzed using MaGECK^43^ and a third screen performed with 10 µM DHA was analyzed separately. The gRNAs against eight genes were significantly enriched in DHA-treated populations in at least two of the three high-dose screens (**Supplementary Table 3**). A further 65 genes were significantly enriched in individual screens (**Supplementary Table 4**). Because K13 resistance alleles retain function and have not been recovered by selection in culture, the absence of K13 from our loss-of-function screens was expected^12,13,17,18,20,21^.

**Figure 3.**
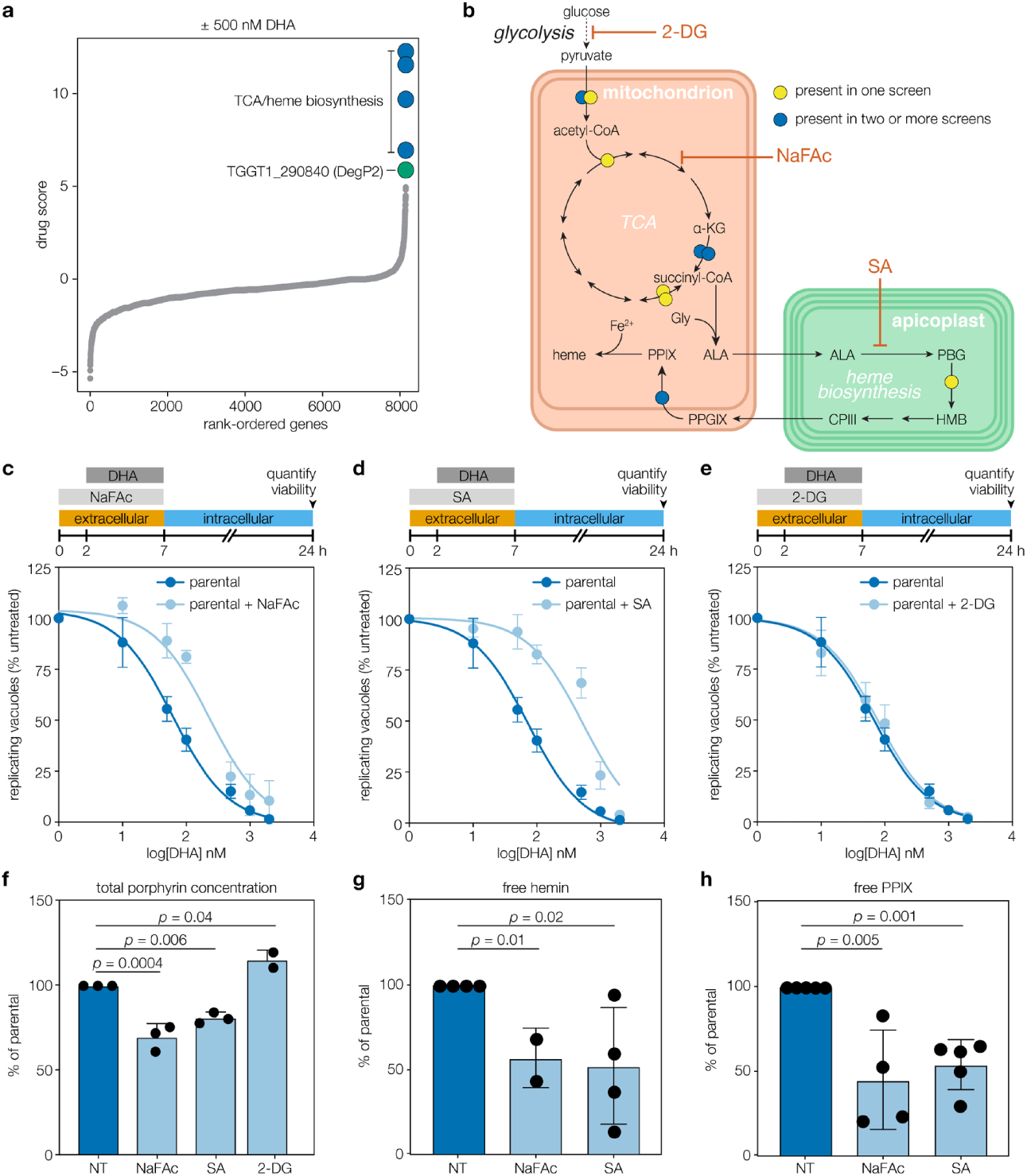
Genome-wide CRISPR screen under high DHA pressure identifies TCA and heme biosynthetic pathways as determinants of DHA sensitivity. **a**, Aggregated drug scores from two independent genome-wide CRISPR screens performed with a lethal dose of 500 nM DHA. Genes with consistently high drug scores are highlighted indicating that their disruption favored parasite survival under otherwise lethal concentrations of DHA. **b**, Diagram of key metabolic pathways highlighting genes with high drug scores in the screens from A and a third screen performed with 10 µM DHA, which was analyzed separately. Predicted localization of enzymes within organelles for the TCA cycle and heme biosynthesis pathways is illustrated. 2-DG, 2-deoxyglucose; α-KG, α-ketoglutarate; ALA, δ-aminolevulinic acid; CPIII, coproporphyrinogen III; Gly, glycine; HMB, hydroxymethylbilane; NaFAc, sodium fluoroacetate; PGB, porphobilinogen; PPIX, protoporphyrin IX; PPGIX, protoporphyrinogen IX; SA, succinyl acetone. Hits found in two or more replicates (blue) or only one replicate (yellow) are indicated. **c**, Pretreatment with 500 mM NaFAc for 2 h increased the IC_50_ of DHA significantly (extra-sum-of-squares F test, *p* < 0.0001). Results are mean for n = 7 or 4 independent experiments with vehicle or NaFAc treatment, respectively. **d**, Pre-treatment with 10 mM SA increased the IC_50_ of DHA significantly (extra-sum-of-squares F test, *p* < 0.0001). Results are mean ± SEM for n = 7 or 5 independent experiments with vehicle or SA treatment, respectively. **e**, Pre-treated with 5 mM 2-DG did not affect the IC_50_ of DHA significantly (extra-sum-of-squares F test, *p* = 0.6). Results are mean ± SEM for n = 7 or 3 independent experiments with vehicle or 2-DG treatment, respectively. **f**, Porphyrin levels (combination of heme and PPIX) in parasites exposed to various compounds, measured by fluorescence and normalized to untreated parasites (NT). Results are mean ± SD for n = 3 independent experiments, each performed in technical duplicate; *p* values from a one-way ANOVA with Tukey’s test. **g**–**h**, Relative abundance of free hemin (g) and free PPIX (h), quantified by LC-MS after treatment with the indicated compound and normalized to untreated parasites (NT). Results are mean ± SD for n = 2–5 independent experiments (indicated by the dots over each column), performed at minimum in technical triplicate; *p* values from a two-way ANOVA.

We performed metabolic pathway analysis on the 73 candidate genes from the screen (65 genes identified in a single replicate and 8 genes identified in multiple replicates) and found that TCA-cycle enzymes were significantly enriched (Bonferroni adjusted *p* value = 1.67 x 10^-5^). The TCA cycle is linked to heme biosynthesis through the production of succinyl-CoA, which is a substrate for ALAS (**Figure 3b**), and among the candidates were two enzymes from the heme biosynthesis pathway: porphobilinogen deaminase (TGGT1_271420) and protoporphyrinogen oxidase (TGGT1_272490). We also retrieved the pyridoxal kinase, which functionalizes the cofactor pyridoxal phosphate, required by several enzymes including δ-aminolevulinic acid synthase (ALAS), which catalyzes the first step of heme biosynthesis^44^. Taken together, our results show that genetically altering heme biosynthesis is an effective way of modulating DHA sensitivity.

### Chemical modulation of heme biosynthesis decreases DHA sensitivity

We investigated whether inhibitors of the TCA cycle (sodium fluoroacetate (NaFAc)^45^) or heme biosynthesis (succinyl acetone (SA)^46^) would recapitulate our genetic results. After pretreating extracellular parasites for 2 h with NaFAc or SA, we treated for a further 5 h with various DHA concentrations, then determined parasite viability. Consistent with our results, both NaFAc and SA significantly increased the IC_50_ of DHA compared to untreated parasites (**Figure 3c–d, Supplementary Table 1**). To ensure that metabolic modulation alone did not impact DHA sensitivity, we blocked glycolysis using 5 mM 2-deoxy glucose (2-DG)^45,47^, as glycolysis is dispensable for TCA activity in the presence of glutamine^48^. Pre-treatment with 2-DG did not change the sensitivity of parasites to DHA (**Figure 3e, Supplementary Table 1**).

To confirm that the inhibitors were having the expected effects on heme production, we quantified total levels of porphyrins—including both heme and its precursor protoporphyrin IX (PPIX)—by measuring porphyrin fluorescence^49^. Porphyrin levels were slightly increased by treatment with 2-DG, whereas treatment with NaFAc or SA significantly depressed porphyrin levels (**Figure 3f**). To distinguish between heme and PPIX—which lacks iron and is not expected to activate DHA^4,50^—we analyzed free and protein-bound porphyrin pools by mass spectrometry (MS). After treatment with either NaFAc or SA, we saw a significant reduction in the levels of both free hemin (the oxidized form of heme found in solution, **Figure 3g**) and PPIX (**Figure 3h**) but not in the protein-bound fractions of these metabolites (**Supplementary Figure 3**). By global polar metabolite profiling, we saw the expected changes in the relevant metabolic pathways following treatment with the inhibitors (**Supplementary Figure 3, Supplementary Table 5**). These results demonstrate that the TCA cycle is required to maintain free porphyrin pools in *T. gondii*, joining the metabolic pathways identified in our screens and confirming that modulation of heme levels can result in decreased DHA sensitivity in apicomplexans.

To test whether increased flux through the heme biosynthesis pathway could hypersensitize parasites to DHA, we supplemented the growth medium with 200 μM ALA, a precursor that stimulates heme biosynthesis in many organisms^51–53^. While ALA treatment increased total porphyrin levels (**Supplementary Figure 3**), MS analysis attributed this increase to PPIX, while free and protein-bound hemin levels remained constant (**Supplementary Figure 3**). The surplus PPIX did not alter DHA sensitivity, consistent with a specific role for heme-bound iron in DHA activation (**Supplementary Figure 3**).

### DegP2 is involved in heme biosynthesis and DHA sensitivity

Guide RNAs targeting TGGT1_290840 were enriched in all replicates of the CRISPR screen performed under a lethal dose of DHA. Previously uncharacterized in *T. gondii*, TGGT1_290840 shares homology with the DegP family of serine proteases (**Figure 3a**). A related *T. gondii* rhoptry protease was recently named DegP^54^, so we named TGGT1_290840 DegP2. DegP2 possesses an identifiable catalytic serine (S569), evident in an alignment with the *Arabidopsis thaliana* plastid protease Deg2^55–57^.

We generated a panel of strains to study the localization and function of DegP2. These included a DegP2 knockout made by replacing the coding sequence with YFP (ΔDegP2; **Figure 4a** and **Supplementary Figure 4**). We complemented this knockout with a C-terminally HA-tagged allele expressed from the *TUB1* promoter (ΔDegP2/DegP2-HA). We also endogenously tagged DegP2 with a C-terminal Ty epitope, then mutated DegP2’s catalytic serine, creating two strains that we refer to as DegP2-Ty and DegP2^S569A^-Ty, respectively (**Figure 4a**). Both the Ty-tagged endogenous DegP2 and ectopically-expressed DegP2-HA colocalized with the mitochondrial marker mtHSP70, and mutating the catalytic serine did not affect this localization (**Figure 4b**).

**Figure 4.**
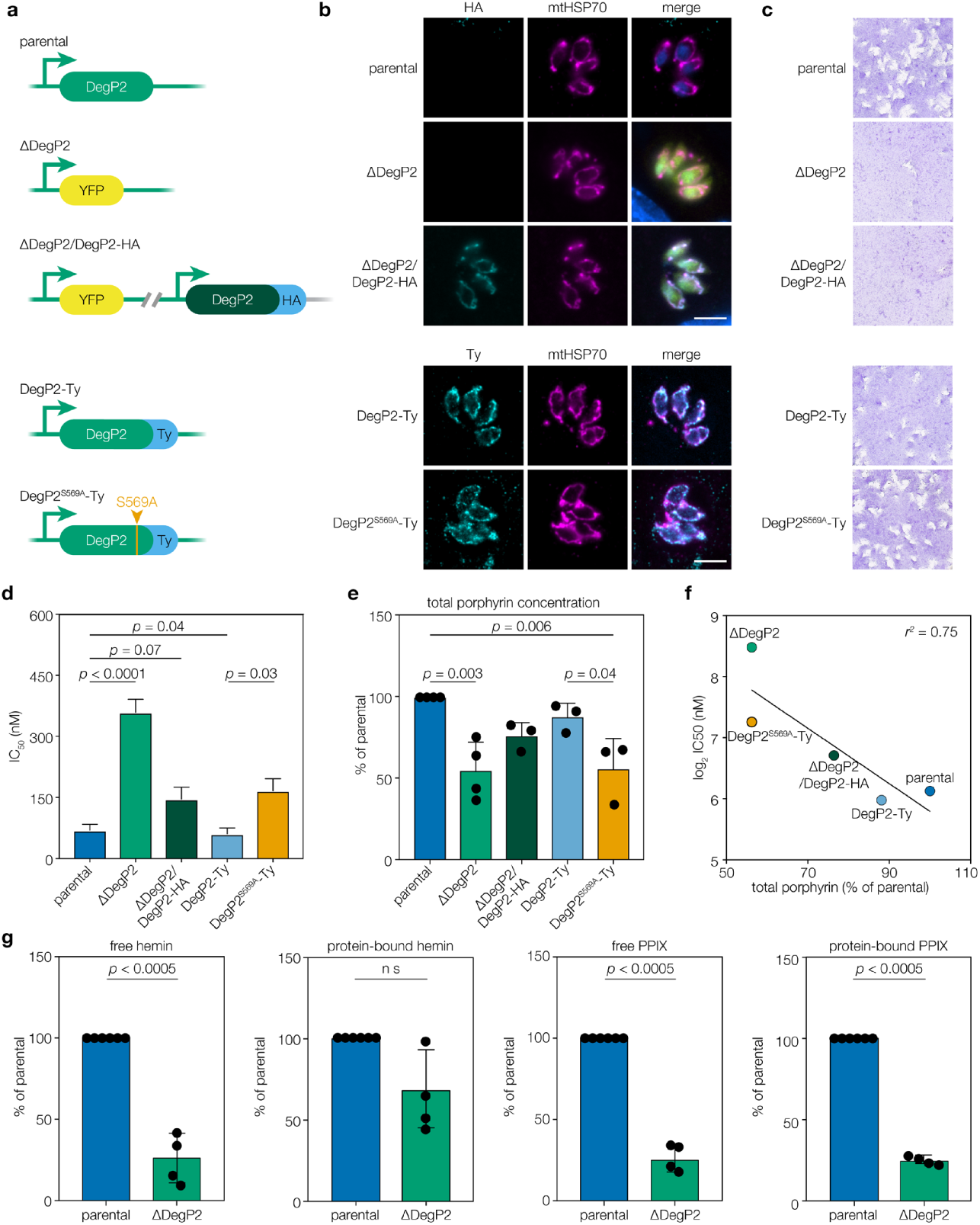
DegP2 has a role in DHA sensitivity and heme biosynthesis. **a**, Diagram of the *DegP2* locus for the various strains used in this work. Regions from the endogenous locus are indicated in light green. The *DegP2* coding sequence was replaced with *YFP* to generate the ΔDegP2 line, which was later complemented with an HA-tagged copy of *DegP2* (dark green) under the regulation of the endogenous 5’ sequences. **b**, DegP2-HA, DegP2-Ty and DegP2^S569A^-Ty co-localized with the mitochondrial marker mtHSP70. Merged image additionally displays YFP (green) for ΔDegP2 strains and DNA stain (blue) for the HA-stained samples, scale bar is 5 μm. **c**, Plaque formation by the various strains. ΔDegP2 and ΔDegP2/DegP2-HA parasites formed smaller plaques than their parental strains, indicating a growth defect. Endogenously Ty-tagged DegP2 (DegP2-Ty) parasites and a derived line bearing a mutation in the protease domain (DegP2^S569A^-Ty) formed plaques normally. **d**, Summary of the DHA IC_50_ values from the DegP2 strains. Results are mean ± SEM for n = 4–7 independent experiments; *p* values from Brown-Forsythe ANOVA controlled for FDR. **e**, Total porphyrin concentrations normalized to the parental strain, showing reduced porphyrin concentrations for both ΔDegP2 and DegP2^S569A^-Ty parasites. Results are mean ± SD for n = 3–4 independent experiments, as indicated by the dots over each column; *p* values are from one-way ANOVA with Tukey’s test performed on the data prior to normalization. **f**, Comparison of DHA IC_50_ (D) with normalised porphyrin levels (E) showing a negative correlation (*r2* = 0.75) in the strains tested. **g**, LC-MS analysis of protein-bound or free hemin and PPIX. Deletion of DegP2 leads to a significant reduction in PPIX and free, but not protein-bound, hemin. Results are mean ± SD for n = 4 independent experiments, each performed at minimum in technical triplicate; *p* values from two-way ANOVA on the data prior to normalization.

Plaque assays were performed with all of the DegP2 lines to assess the effect of the mutations on the lytic cycle of *T. gondii*. Both ΔDegP2 and the complemented strain (ΔDegP2/DegP2-HA) had significant growth defects, producing small plaques, although they could be maintained in culture. By contrast, DegP2-Ty and DegP2^S569A^-Ty lines formed plaques similar to the parental strain (**Figure 4c**). Since the growth defect of ΔDegP2 could not be complemented, despite restoration of other DegP2 functions, the knockout may harbor alterations that extend beyond the activity of the protease. We therefore consider the overlap between ΔDegP2 and DegP2^S569A^-Ty to best represent the pathways impacted by DegP2 activity.

To investigate whether loss of catalytic activity or deletion of DegP2 altered DHA sensitivity, we determined the DHA IC_50_ of each strain in our panel of mutants (**Figure 4d, Supplementary Figure 4**). ΔDegP2 parasites showed reduced DHA sensitivity compared to parental, while intermediate responses were observed for the complemented strain and DegP2^S569A^-Ty (**Figure 4d, Supplementary Table 1**). Based on its localization and the relationship to heme that was shared by many of our other screening hits, we predicted that disrupting DegP2 might decrease parasite heme levels. We found that either deleting DegP2 (ΔDegP2) or ablating its protease activity (DegP2^S569A^-Ty) significantly reduced porphyrin levels (**Figure 4e**), which was inversely correlated with DHA sensitivity (**Figure 4f**). MS showed that free hemin accounted for some of this change in both the ΔDegP2 and DegP2^S569A^-Ty lines, (**Figure 4g** and **Supplementary Figure 4**), along with decreased free and protein-bound PPIX (**Figure 4g**). This indicates that DegP2 modulates heme metabolism in *T. gondii.*

The decreased heme levels in ΔDegP2 prompted the examination of the other mutants that modulate DHA sensitivity. ΔTmem14c and K13^C627Y^ parasites displayed similar levels of porphyrins and free heme when analyzed by fluorescence or MS (**Supplementary Figure 2**). Analysis of polar metabolitesn in the K13^C627Y^ strain did reveal a significant change (FDR < 0.05) in the metabolic pathway for alanine, aspartate, and glutamate production. TCA cycle was also affected (FDR < 0.1) with several intermediates slightly lower than in the control (**Supplementary Figure 2, Supplementary Table 6**). Unlike for NaFAc (**Supplementary Figure 3**) or ΔDegP2 (discussed below), this effect does not indicate an asymmetry in the flux through the pathway. Our results therefore suggest that in *T. gondii*, ΔTmem14c and K13^C627Y^ mutations affect DHA sensitivity independently from the steady-state heme pools, although we cannot rule out changes in subcellular localization of heme, as may be expected from the predicted function of Tmem14c as a porphyrin transporter^42^.

To determine whether heme biosynthesis is active in the absence of DegP2, we supplemented ΔDegP2 parasites with ALA and measured downstream metabolites by MS. ALA supplementation increased levels of free and protein-bound PPIX, indicating that there is no intrinsic deficiency in the ability of ΔDegP2 parasites to synthesize PPIX. However, as observed with parental parasites, hemin levels did not increase when the mutants were supplemented with ALA (**Supplementary Figure 4**), demonstrating that recovery of PPIX is not sufficient to reverse the defect seen in heme levels in the ΔDegP2 line.

### Absence of DegP2 leads to subtle changes in mitochondrial homeostasis

To further examine the function of DegP2, we compared gene expression in ΔDegP2 and parental parasites. By RNAseq, 20 genes were significantly down-regulated and 6 were up-regulated in the ΔDegP2 line (*p* adj < 0.05). Three of these up-regulated genes are encoded by the mitochondrial genome in *P. falciparum* ^58^, and may similarly be encoded by the *T. gondii* mitochondrial genome for which a complete sequence is not available. The proteins encoded by these upregulated transcripts (*cox1*, TGGT1_408770; *cox3*, TGGT1_365020; *cyt b*, TGGT1_409830) function in the electron transport chain (ETC; **Figure 5a** and **Supplementary Table 7**). Intriguingly, manipulation of heme levels has been shown to affect mitochondrial transcription in other organisms^59,60^. We also observed changes in the abundance of TCA-cycle metabolites including pyruvate, glutamine, malate, succinate and GABA when comparing ΔDegP2 to the parental line (**Figure 5b** and **Supplementary Table 8**).

**Figure 5.**
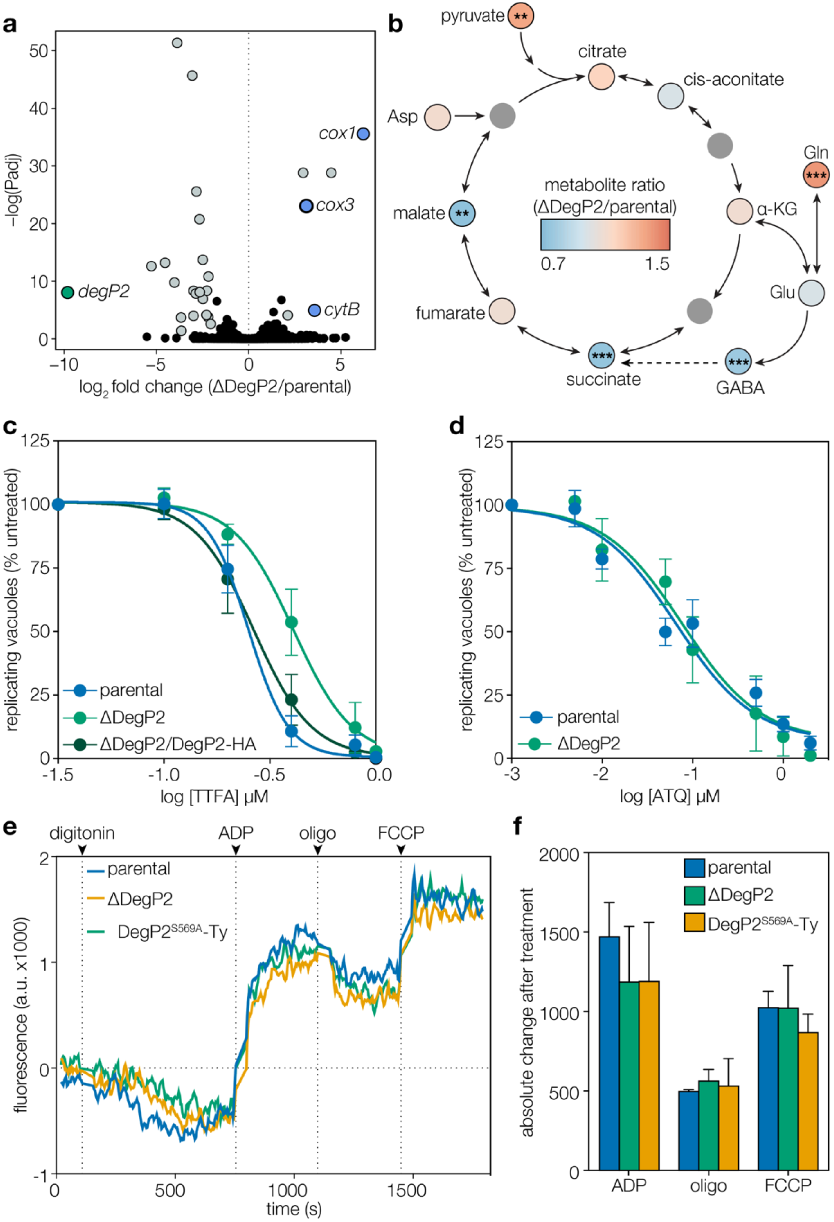
Deletion of DegP2 alters the TCA cycle and the electron transport chain. **a**, Volcano plot showing fold change of transcripts between parental and ΔDegP2 parasites. Genes with adjusted *p* values (Padj) < 0.05 are highlighted in grey. Three cytochrome genes (blue) were significantly upregulated upon deletion of DegP2. As expected, transcripts for DegP2 were significantly downregulated in the knockout (green). **b**, Summary of changes in metabolites between parental and ΔDegP2 parasites in the TCA cycle and closely related pathways. Full results can be found in **Supplementary Table 8.** Asp, aspartate; α-KG, α-ketoglutarate; Gln, glutamine; Glu, glutamate. Asterisks indicate significant change from parental, **\*** *p* < 0.05, ** *p* < 0.005, *** *p* < 0.0005, by two-way ANOVA. **c**, Dose-response curve for parasites treated with the complex II inhibitor TTFA. Results are mean ± SEM for n = 4 or 3 independent experiments, for the parental and ΔDegP2 or ΔDegP2/DegP2-HA strains, respectively. ΔDegP2 is significantly more resistant to TTFA compared to the parental; *p* < 0.0001 from extra-sum-of-squares F test. **d**, Dose-response curve for parasites treated for 5 h with increasing concentrations of atovaquone (ATQ). Results are mean ± SEM for n = 5 or 3 independent experiments for the parental or ΔDegP2 strains, respectively. Both strains displayed similar sensitivity to ATQ; *p* = 0.59 from extra-sum-of-squares F test. **e**–**f**, Mitochondrial polarization was monitored using Safranin O fluorescence in digitonin-permeabilized cells. Parasite mitochondria were allowed to polarize in the presence of succinate before adding ADP, which triggers depolarization as complex V uses the proton-motive force to synthesize ATP. Oligomycin (oligo) inhibits complex V causing repolarization, while FCCP completely depolarizes the mitochondria. Traces from a representative experiment are shown (e). Changes in fluorescence were consistent after each drug treatment for all strains tested (F). Results are mean ± SD for n = 3 independent experiments.

To determine whether DegP2 influences ETC activity, we measured sensitivity to the Complex II inhibitor Thenoyltrifluoroacetone (TTFA)^61,62^, and the cytochrome *b* inhibitor atovaquone (ATQ)^63,64^. The parental and complemented lines showed similar responses to TTFA, while ΔDegP2 parasites were significantly more resistant (**Figure 5c, Supplementary Table 1**). By contrast, the IC_50_ of ATQ was unchanged between ΔDegP2 and the parental line (**Figure 5d, Supplementary Table 1**). We also compared mitochondrial polarization using Safranin O: a self-quenching fluorescent dye that concentrates in polarized mitochondria^65,66^. Mitochondrial polarization was achieved in semi-permeabilized parasites using succinate—a substrate for Complex II. Loss of DegP2 did not affect the mitochondrial response to ADP, oligomycin, or Carbonyl cyanide 4-(trifluoromethoxy)phenylhydrazone (FCCP; **Figure 5e–f**). Comparable mitochondrial polarization was also observed by MitoTracker-loading (**Supplementary Figure 4f**). These results indicate a subtle and non-uniform dysregulation of the ETC in the DegP2 mutant.

### SA treatment reduces DHA sensitivity in *P. falciparum*

To determine if free-heme levels correlate with *P. falciparum*’s sensitivity to DHA in the heme-rich environment of the red blood cell, we used the ring-stage survival assay (RSA) to examine the effect of blocking heme biosynthesis with SA on the sensitivity of parasites to DHA. The Cambodian isolate Cam3.II carrying the R539T mutation in K13, as well as two isogenic lines that were either reverted back to the K13 wild-type sequence (Cam3.II WT) or genetically modified to carry the K13^C580Y^ allele^13^, were tested for the effect of SA on DHA sensitivity. SA pretreatment significantly decreased the DHA sensitivity of Cam3.II carrying a wild-type allele of K13; however, in parasites carrying either K13^C580Y^ or K13^R538T^ mutant alleles, SA treatment did not increase DHA survival further (**Figure 6a**).

**Figure 6.**
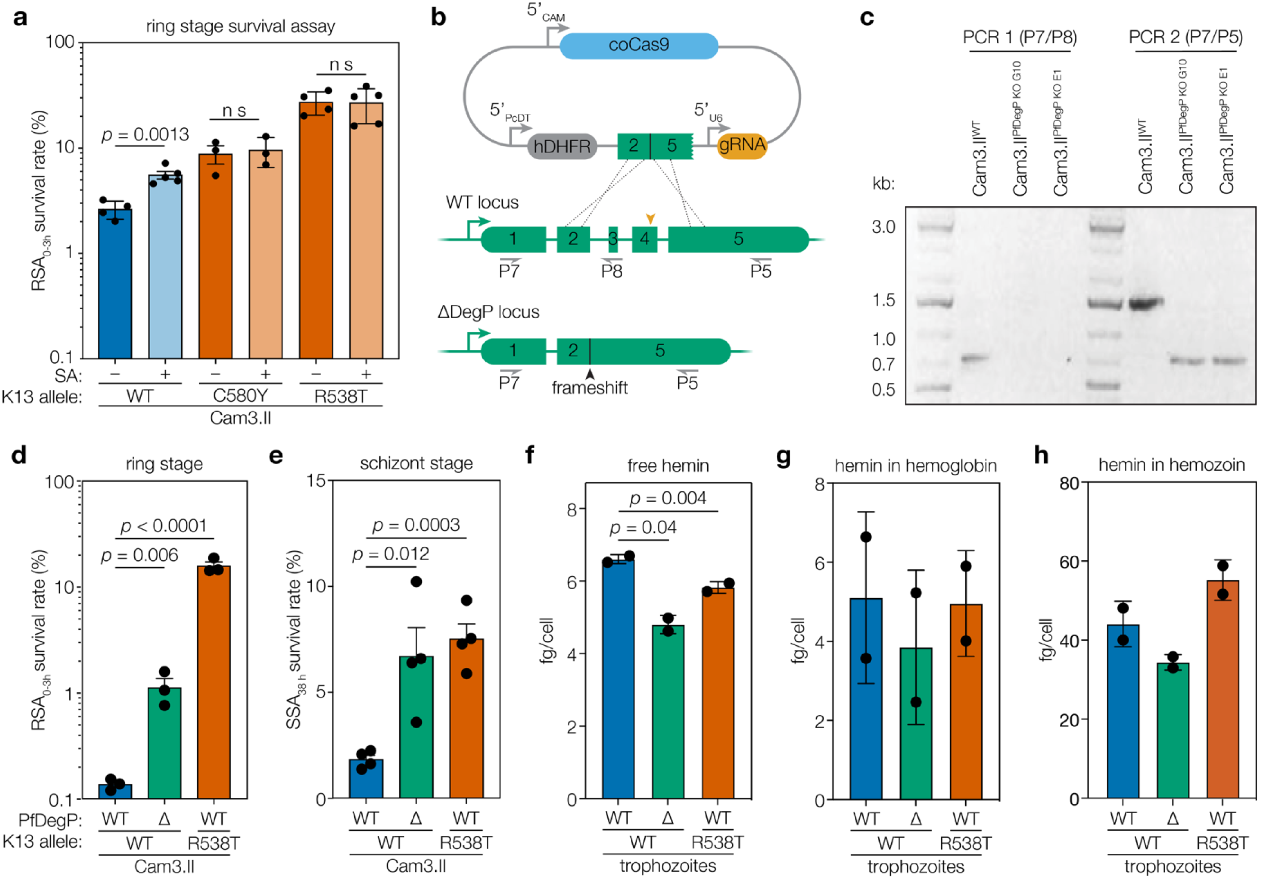
SA treatment and the DegP2 ortholog similarly affect DHA sensitivity in *P. falciparum*. **a**, Ring-stage survival assay, following a 12 h pretreatment of schizonts with 200 μM SA or a vehicle control. Percentage of early ring-stage *P. falciparum* (0 to 3 h after invasion) surviving a 4 h pulse of 700 nM DHA, assessed by flow cytometry 66 h later. Results are mean ± SEM for n = 4 independent experiments; *p* values are from unpaired *t* test. **b**, Diagram showing strategy to knock out PfDegP in the Cam3. II^WT^ background. **c**, PCR confirmation of successful KO og PfDeg2, PCR 1 (using primers P7 and 8) confirming loss of exons 3 and 4 in the two DegP KO clones used in this study. PCR 2 (using primers P7 and 5) demonstrates a shorter product upon successful deletion of two exons. **d**, Ring-stage survival assay, performed as above, using the Cam3.II^WT^, Cam3.II^ΔPfDegP^ and the K13 mutant Cam3.II^R539T^. Results are mean ± SEM for n = 4 independent replicates; *p* values are from unpaired *t* test. **e**, Schizont-stage survival assay following a 4 h pulse of 700 nM DHA. Results are mean ± SEM for n = 4 independent replicates; *p* values are from unpaired *t* test. **f**–**h**, Heme measured from the free (f), hemoglobin-associated (g), and hemozoin pools from trophozoites for various strains. Heme concentrations were calibrated to a standard curve and normalized to the number of parasites per sample to calculate fg/cell. Results are mean ± SD for n = 2 independent replicates; *p* values from one-way ANOVA.

### PfDegP deletion alters heme concentrations and reduces DHA sensitivity during the erythrocytic cycle

An ortholog of DegP2 is readily identifiable in *P. falciparum*. To investigate its role in artemisinin sensitivity, the DegP2 homolog (PF3D7_0807700) was knocked out (**Figure 6b–c**) in the Cam3.II strain carrying the wild-type allele of K13^13^. We compared the DHA sensitivity of Cam3.IIΔPfDegP parasites to the parental strain and the K13^R539T^ strain. DHA sensitivity was assessed by RSA and by a similar assay that measures sensitivity during the schizont stage (SSA). The K13^R539T^ mutation decreased DHA sensitivity in the RSA (**Figure 6d**), as previously demonstrated^13^, and in the SSA (**Figure 6e**). Deleting *PfDegP* also decreased the DHA sensitivity of both ring-stage and schizont-stage parasites (**Figure 6d–e**), although the effect on ring-stage parasites was less pronounced than that of the K13 mutation.

To test whether the changes in DHA sensitivity were correlated with changes in heme concentrations, we fractionated infected cells and measured free heme, as well as heme associated with hemoglobin and hemozoin^67,68^. Trophozoites from either the *PfDegP* knockout or those carrying the K13^R539T^ mutation had significantly lower levels of free heme (**Figure 6f**), while heme associated with hemoglobin or hemozoin remained unchanged (**Figure 6g–h**). The effect of the mutation on trophozoites points to a link between DHA sensitivity and the availability of free heme within the parasite. Taken together, our results demonstrate that screens in *T. gondii* can identify resistance alleles that act through mechanisms conserved across the phylum.

## DISCUSSION

Our results point to a central role for heme biosynthesis in the sensitivity of apicomplexan parasites to DHA. Through genetic and chemical perturbations, we demonstrate that the levels of heme available for the predicted activation of artemisinin^4,22,52,69^ can be modulated sufficiently to affect sensitivity. Our results help identify new components of the heme biosynthetic pathway, and inform additional pathways that should be taken into account when designing artemisinin combination therapies.

Several observations are consistent with a limiting role for heme in artemisinin sensitivity in parasites. *Plasmodium* spp. and *T. gondii* are more sensitive to artemisinins than mammalian cells, where labile heme is bound to chaperones and free heme is often undetectable^70,71^. We also found that levels of free heme were a better indicator of reduced DHA sensitivity than the protein-bound pool in *T. gondii*. Relatedly, the apicomplexan parasite *Cryptosporidium parvum*—which lacks genes of the heme biosynthetic pathway^72,73^—shows little response to artemisinin^74^. Furthermore, the availability of free heme varies among *Plasmodium* stages in concert with sensitivity to artemisinin^75^. Trophozoites, which have high levels of free heme from degrading hemoglobin, are more sensitive to artemisinins than ring or liver stages, which are thought to rely predominantly on their endogenous heme biosynthetic pathways^38,76,77^. The implication of endocytic machinery in artemisinin sensitivity^18,20,21^ could also be related to reduced hemoglobin uptake and digestion, which would limit or delay the availability of free heme.

CRISPR-based screens can identify sensitizing mutations, which may help design combinatorial therapies. Screening for sensitivity to a sublethal dose of DHA, we identified an ortholog of Tmem14c, which is thought to transport porphyrins across the inner mitochondrial membrane of mammalian cells^42^. The precise role of Tmem14c in *T. gondii* still needs to be determined; however, the increased sensitivity of intracellular knockouts implies its function may be more critical during replication. Dramatic changes in *T. gondii* metabolism have been observed between intracellular and extracellular parasites^45,78^. The decreased heme biosynthesis measured following SA or NaFAc treatment argues for active biosynthesis in extracellular parasites; however, replicating parasites likely demand an increased rate of heme biosynthesis, which may explain why ΔTmem14c parasites were more sensitive to DHA during this period of their lytic cycle. Technical limitations precluded reliably measuring heme in replicating parasites, but the predicted role of Tmem14c as a porphyrin transporter leads us to propose that changes in heme availability, such as its accumulation in the mitochondrion, may influence the sensitivity of parasites to DHA^71,79^.

Homologous mutations in K13 decrease the sensitivity of both *T. gondii* and *P. falciparum* to DHA. However, the evidence linking K13 and heme is inconclusive. Extracellular K13^C627Y^ *T. gondii* have similar heme concentrations to those found in the parental strain. By contrast, we observed lower heme concentrations in *P. falciparum* K13^R538T^ trophozoites. Moreover, disrupting heme biosynthesis in *P. falciparum* using the ALAD inhibitor SA reduced DHA sensitivity in K13^WT^ parasites, but did not further enhance survival of strains bearing the resistance alleles. As with Tmem14c, heme compartmentalization not captured by our bulk measures could affect DHA sensitivity; e.g., K13 mutations may affect mitochondrial or vacuolar free heme without affecting total heme concentrations. The reduction in DHA sensitivity caused by SA treatment of *P. falciparum* rings—which derive their heme mainly from mitochondrial pathways^80^—supports the notion that DHA activation in ring stages depends on mitochondrial heme^4^.

Screening for mutants with decreased compound sensitivity identified the serine protease DegP2, knockout of which contributed to the survival of *T. gondii* and *P. falciparum* under DHA treatment. In both species, the activity of DegP2 (*Pf*DegP) could be linked to decreased free heme concentrations. Deg-family proteins play roles in the folding, maintenance, and turnover of integral membrane proteins, such as those involved in the electron transport chain^81–83^. PfDegP complements its ortholog in *E. coli*, implying functional conservation^84^. Maintenance of mitochondrial membrane proteins could explain the observed impact of DegP2 on mitochondrial pathways. Increased expression of ETC genes could therefore reflect the successful compensatory response to reduced heme levels in ΔDegP2, as observed in yeast and mammals^85–87^. Our results argue for a conserved role for DegP2 across *T. gondii* and *P. falciparum* in mitochondrial activity and DHA sensitivity.

We show that activation of DHA by free heme provides a critical bottleneck, and that heme deficient *T. gondii* and *P. falciparum* escape the short-lived, intense drug pressure caused by artemisinin. Severely compromised mutants in TCA-cycle enzymes^32^ were enriched when we performed a CRISPR screen using a lethal dose of DHA, demonstrating that this approach can reveal mutations that protect against drug pressure, even when such mutations are highly detrimental in other contexts. This highlights the power of genome-wide CRISPR screens in identifying mechanisms of drug activity and resistance, and points to the role they can play in developing novel therapeutics.

## ACKNOWLEDGEMENTS

We would like to thank the MIT Genome Core and the Whitehead Metabolomics Core for technical assistance. Emily Shortt and Eric Spooner assisted with proteomics. Sheena Vasquez and Jade Bath helped adapt the porphyrin and Safranin O assays, and Bastien Christ assisted with mass spectrometry– based heme measurements. George Bell provided statistics advice. This study was supported by a Sir Henry Wellcome fellowship to C.R.H. (103972/Z/14/Z), an NIH R01 to D.A.F. (AI109023), and an NIH Director’s Early Independence Award (1DP5OD017892) and a Mathers Foundation grant (MF-1706-00164) to S.L.

## MATERIALS & METHODS

### *T. gondii* maintenance and strain construction

All *T. gondii* strains were derived from RH Δku80 parasites^88^. *T. gondii* tachyzoites were grown in human foreskin fibroblasts (HFFs) maintained at 37 °C with 5% CO_2_ in D3 [Dulbecco’s modified Eagle’s medium (DMEM, Life Technologies, 11965118) supplemented with 3% heat-inactivated fetal bovine serum and 10 μg/mL gentamicin (Sigma Aldrich, G1272)]. When appropriate, chloramphenicol (Sigma Aldrich, C3175) was used at 40 μM and pyrimethamine (Sigma Aldrich, 46706) at 3 μM.

Parasite lines were constructed using CRISPR/Cas9-mediated gene editing as described previously^89^. The Cas9-encoding plasmid pU6-Universal is available from Addgene (#52694). Gibson Assembly^90^ was used to clone gRNAs into the BsaI sites in pU6-Universal using homology arms of approximately 20 base pairs. Epitope tags and point mutations were introduced by transfecting repair oligos, bearing 40 base pairs of homology with the genome on each end, along with CRISPR machinery. Such repair oligos were often created by heating oligonucleotides encoding complementary single-stranded DNA to 99 °C for 2 minutes in duplex buffer (100 mM potassium acetate, 30 mM HEPES, pH 7.5), then allowing DNA to cool slowly to room temperature over the course of several hours.

K13^C627Y^ mutant parasites were constructed by transfecting pU6-Universal encoding the protospacer TTGCTCCTCTCACCACTCCG (oligos P1 and P2 in **Supplementary Table 9**) into Δku80 parasites along with a repair oligo constructed by hybridizing the oligos P3 and P4. Converting the TGC that encodes C627 to TAC eliminated an *ItaI* site, and restriction digests were used to confirm this mutation. We also introduced a silent CGG to CGA mutation at R667 to eliminate the protospacer adjacent motif (PAM) and prevent re-cleavage by Cas9 after the initial mutation. Clonal K13^C627Y^ lines were isolated, and the mutation was confirmed by sequencing the locus using primers P5 and P6.

ΔDegP2 parasites were constructed by co-transfecting Δku80 parasites with two copies of pU6-Universal encoding gRNAs with the sequences GCAGTCCCCAGCATGGTCGG (P7 and P8 in **Supplementary Table 9**) and GCGCTCACAAGACCTCGCTGG (P9 and P10) to cleave the 5’ and 3’ ends of the ORF respectively. These cuts were repaired using a template encoding YFP flanked by 40 nucleotides of homology to the 5’ and 3’ UTRs of DegP2, constructed by amplifying the fluorescence marker with P11 and P12. YFP-positive parasites were selected by FACS, and the insertion was confirmed by PCR and Sanger sequencing using primers P13 and P14. Later, YFP was mutated by transfecting parasites with pU6-Universal carrying the guide sequence GGTGCCCATCCTGGTCGAGC (P46 and P47) and a repair template constructed by hybridizing the primers P44 and P45, which introduces three stop codons near its 5’ end. YFP-negative parasites were selected by FACS and a clonal population was isolated using limiting dilution.

DegP2-Ty parasites were constructed by transfecting pU6-Universal carrying the gRNA sequence GCGCTCACAAGACCTCGCTGG, together with hybridized P15 and P16, into the Δku80 strain. Clonal lines were isolated using limiting dilution, and those carrying the Ty tag were identified using immunofluorescence microscopy^91^.

ΔDegP2 was complemented by integrating the plasmid pDegP2-HA at the intergenic region at position 1487300 on chromosome VI. pDegP2-HA was constructed by amplifying DegP2 from cDNA using the primers P17 and P18 and assembling it with a portion of pDsRed^92^ that was amplified using the primers P19 and P20. A 200 nucleotide region surrounding the integration locus, with an *NheI* site at its center, was cloned into the *PstI* site of the resulting plasmid. We co-transfected *NheI-*linearized pDegP2-HA with a Cas9-expressing plasmid carrying a gRNA with the sequence GCCGTTCTGTCTCACGATGC, then selected for integrants using 40 μM chloramphenicol. Clonal populations carried DegP2-HA were confirmed using immunofluorescence microscopy with an anti-HA antibody (16B12, Biolegands).

DegP2^S569A^-Ty parasites were constructed by transfecting DegP2-Ty parasites with pU6-Universal encoding a protospacer with the sequence GCCATTAATCCTGGCAACAG (oligos P21 and P22) and a repair oligo constructed by annealing P23 and P24. Positive clones were selected based on destruction of a *HpyCH4III* restriction site and confirmed by PCR and Sanger sequencing using primers P23 and P24.

ΔTmem14c parasites were constructed by co-transfecting Δku80 parasites with two copies of pU6-Universal encoding gRNAs with the sequence TCGGATTGCTATCTGACCAA (oligos P27 and P28) and ACGTCTGATGCCAAGGCGAT (primers P29 and P30), to cleave the 5’ and 3’ ends of the ORF, respectively. These cuts were repaired using a PCR product encoding a *TUB1* promotor, mNeonGreen coding sequence, and the *SAG1* 3’ UTR. This sequence was amplified using the primers P31 and P32. mNeonGreen-positive parasites were selected by FACS, and the insertion was confirmed by PCR and Sanger sequencing using the primers P33 and P34, P35 and P36 and P37 and P38.

Tmem14c-Ty parasites were created by randomly integrating a plasmid containing Tmem14c-ty into RH parasites. Through PCR and sanger sequencing of cDNA, the gene model on ToxoDB was found to be incorrect. The stop codon is instead found at the start of exon 2, creating a product of 330 bp. This product was amplified from cDNA using the primers P36 and P39 and assembled into the pSAG1-GCaMP5^93^ backbone in which the chloramphenicol-resistance cassette was replaced with *DHFR*, in frame with a Ty tag between the *SAG1* promoter and 3’-UTR. Parasites were selected using pyrimethamine, cloned by limiting dilution and positive clones selected by immunofluorescence microscopy for the Ty tag.

### Pooled genome wide screens

Genome-wide CRISPR screens were performed based on the previously published method^31^. Briefly, 500 mg of gRNA library linearized with *AseI* was transfected into approximately 4 × 10^8^ Cas9-expressing parasites^40^ divided between 10 individual cuvettes. Parasites were allowed to infect 10 x 15 cm^2^ plates of confluent HFFs and the medium changed 24 h post infection to contain pyrimethamine and 10 μg/ mL DNaseI. Parasites were passaged upon lysis (between 48–72 h post transfection) and selection continued for three passages. At this point, parasites with integrated gRNA plasmids were split into two pools and either treated with indicated concentrations of DHA or left untreated. Untreated parasites were passaged onto fresh HFFs every two days while the medium on treated parasites was changed for fresh DHA at the same time point. After a further three passages of the untreated population, parasites were collected, counted and gDNA was extracted using a DNeasy blood and tissue kit (Qiagen). When parasites were not limiting (e.g., in the untreated sample) DNA was extracted from 1 x 10^8^ parasites. When DHA was used for positive selection, DNA was extracted from all recovered parasites. Integrated gRNA constructs were amplified from 500 ng of extracted DNA using the primers P40 and P41 in two independent reactions and pooled. The resulting libraries were sequenced on a HiSeq 2500 (Illumina) with single-end reads using primers P42 and P43.

### Statistical analysis

Sequencing reads were aligned to the gRNA library. The abundance of each gRNA was calculated and normalized to the total number of aligned reads. Guides that were not found were assigned a pseudo-count corresponding to 90% of the lowest value in that sample. Only guides whose abundance was above the 5^th^ percentile in the original plasmid preparation of the gRNA library were taken into account for subsequent analyses. After normalization to guide abundance in the original library, the log_2_-fold change between DHA treated and untreated samples was calculated for each gRNA. The drug score for each gene was calculated as the mean log_2_ fold change for the top five most abundant guides in the library, which minimized the effect of stochastic losses and decreased the error between biological replicates. To determine the fitness effect of each guide under DHA across biological replicates we made use of the Model-based Analysis of Genome-wide CRISPR/Cas9 Knockout (MAGeCK) algorithm^43^. Hits were selected based on FDR of less than 0.1 in at least one biological replicate. Metabolic pathways analysis on hits was performed using ToxoDB^94^ against KEGG and MetaCyc pathways and the Bonferroni adjusted *p* value reported.

### Compounds used with *T. gondii*

Dihydroartemisinin (DHA; VWR, TCD3793) was prepared at 10 mM in DMSO as single use aliquots and used at the indicated concentration. Hemin (Sigma Aldrich, H9039) was prepared at 10 mM in 0.5 M NaOH. 2-Deoxyglucose (Sigma Aldrich, D6134) was prepared fresh at 10 mM in D3 and used at a final concentration of 5 mM. Succinyl acetone (SA; Sigma Aldrich, D1415) was prepared in PBS as 200 mM stocks and used at a final concentration of 10 mM. ALA (Sigma Aldrich, A3785) was prepared fresh in PBS at 200 mM and used at a final concentration of 200 μM. Sodium fluoroacetate (NaFAc; Fisher Scientific, ICN201080) was prepared at 1 M in water and used at a final concentration 500 µM. Atovaquone (Sigma Aldrich, A7986) was prepared at 27 mM in DMSO and used at the indicated concentration. Thenoyltrifluoroacetone (TTFA; Sigma Aldrich, T27006) was prepared as 10 mM stocks in DMSO and used at the indicated concentrations. MitoTracker Deep Red FM (Life Technologies, M22426) was used at a final concentration of 50 nM. Oligomycin (EMD Millipore, 495455) was used at a final concentration of 20 μM. Pyrimethamine (Sigma Aldrich, 46706) was prepared in ethanol at 10mM. ADP (A2754-100MG) was prepared at 100 mM in water. FCCP (Sigma-Aldrich C2920) was prepared at 50 mM in DMSO.

### Lytic assay

50,000 parasites/well were spun down onto HFFs grown in 96-well plates. After one or 24 hours, the medium was removed and replaced with medium containing vehicle or drug and parasites were incubated for the indicated time before the wells were washed, the medium replaced, and the monolayers incubated until 72 h p.i. Monolayers were rinsed with PBS, fixed in 95% ethanol or 100% ice-cold methanol for 10 minutes and stained with crystal violet (Sigma, C6158) stain (2% crystal violet, 0.8% ammonium oxalate, 20% ethanol) for 5 minutes before washing excess dye with water and allowing to dry. Absorbance of wells was read at 570 nm and normalised to drug treated, uninfected wells. Experiment performed in triplicate and results are mean of at least 4 independent experiments.

### Pulsed treatment assay

Extracellular parasites were incubated in a total volume of 100 μl D3 with DHA for the indicated time, spun down and resuspended in 600 μl of pre-warmed D3 before aliquoting 200 μl of parasites to HFF monolayers on a clear-bottomed plate and incubating for 24 h. Wells were then fixed in 4% formaldehyde for 10 minutes and stained using rabbit anti-PCNA and detected using anti-rabbit Alexa 549. Images were acquired using a Cytation 3 high content imager and the number of vacuoles and parasite nuclei per vacuole were quantified using an automated ImageJ macro. Vacuoles of two or more parasites were considered to be alive, and the number of such vacuoles in drug-treated wells was normalized to the number in untreated wells to quantify the lethality of drug treatment.

### Oxalic acid assay

Based on published methods^49^, approximately 5 x 10^7^ freshly-egressed parasites were passed through a 3 μm filter, washed once in 10 mL of PBS, then pelleted and resuspended in 50 μl of H_2_O and flash frozen in liquid nitrogen. 20 μl of parasite lysate was then mixed with 200 μl of pre-warmed 1.5 M oxalic acid (Sigma Aldrich, 75688) and incubated in a thermocycler at 99 °C for 30 minutes. The total volume was then transferred to a clear bottomed plate and fluorescence was detected at 662 nm following excitation at 400 nm. Porphyrin levels were calculated from a standard curve prepared as above using hemin. Total porphyrin was normalised to protein concentration, as calculated by Bradford assay (BioRad, 5000201). Experiments were performed in duplicate and results are representative of at least three independent experiments.

### Immunofluorescence microscopy

Infected cells were fixed in 4% formaldehyde for 10 minutes, then permeabilized with 0.25% Triton X-100 for 8 minutes. Staining was performed with mouse anti-Ty^91^, rabbit anti-mtHSP70^95^, and mouse anti-HA (Biolegends 901513) and detected with Alexa-Fluor-labeled secondary antibodies. Nuclei were stained with Hoechst 33258 (Santa Cruz, SC-394039) and coverslips were mounted in Prolong Diamond (Thermo Fisher, P36961). Images were acquired using an Eclipse Ti epifluorescence microscope (Nikon) using the NIS elements imaging software. FIJI^96^ was used for image analysis and Adobe Photoshop for image processing.

### Plaque formation

500 parasites per well were used to analyze the effect of drug treatment or gene deletion over the course of 8 days. The monolayers were then rinsed with PBS, fixed in 95% ethanol or 100% ice-cold methanol for 10 minutes and stained with crystal violet (2% crystal violet, 0.8% ammonium oxalate, 20% ethanol) for 5 minutes.

### MitoTracker Analysis of ΔDegP2

Parasites were suspended at 1 x 10^7^ parasites per mL in D3 with or without 30 μM oligomycin. After 15 minutes at 37 °C, the samples were mixed 2:1 with MitoTracker in D3, bringing the final concentration of oligomycin to 20 μM and MitoTracker to 12.5 nM. Parasites were incubated at 37 °C for an additional 30 minutes, then pelleted and resuspended in Ringer’s solution (115 mM NaCl, 3 mM KCl, 2 mM CaCl_2_, 1 mM MgCl_2_, 3 mM NaH_2_PO_4_, 10 mM HEPES, 10 mM glucose, 1% FBS), before analysis by flow cytometry.

### Transcriptomics

Intracellular parasites were released from HFFs by scraping infected monolayers and then passing the cell milieu through 27-gauge needles. Duplicate samples, each containing 2.5 x 10^6^ ΔDegP2 or parental parasites, were separated from host-cell debris by passing them through 3 μm filters and washed once in PBS. RNA was extracted using an RNeasy kit (Qiagen, catalogue # 74136). Residual DNA was removed using a DNA removal kit (ThermoFisher Scientific catalogue # AM1906), then RNA was concentrated using an RNeasy MinElute clean-up kit from Qiagen (catalogue # 74204). Samples were then prepared using Kapa mRNA hyperprep and analyzed on a HiSeq 2000 using single-end 40 base pair reads with six nucleotide indices. Reads were aligned to the *T. gondii* genome (release 36) using STAR aligner^97^, and changes in transcript frequency were analyzed using DESeq2^98^.

### Mitochondrial polarization assay

Parasites were separated from cell debris by passing them through 3 μm filters, then washed thoroughly in fluorobrite medium (Gibco, A18967-01) and resuspended at 1 x 10^9^ parasites per mL in fluorobrite. Parasites were diluted to 5 x 10^7^ parasites per mL in reaction buffer (125 mM sucrose, 6 mM KCl, 10 mM HEPES-KOH, pH 7.2, 1 mM MgCl_2_, 2.5 mM potassium phosphate). Safranin O (Sigma Aldrich, S-2255) and succinate (Sigma Aldrich, W327700-SAMPLE-K) were added to final concentrations of 1.25 μM and 1 mM, respectively. For each strain, 160 μl of parasite suspension was added to each of 6 wells in a 96 well plate (Perkin Elmer, 6055300). Parasites fluorescence was recorded every four seconds for 100 seconds using a Cytation 3 high-content imager at an excitation wavelength of 495 nm and an emission wavelength of 586 nm. Ten microliters of 600 μM digitonin in cold reaction buffer was added to each well, and the fluorescence was recorded every four seconds for an additional ten minutes. Three wells of parasites per strain were then subjected to treatment with 10 μl of three drugs sequentially: 200 μM ADP, 2 μg/mL oligomycin, and 20 μM FCCP, all prepared in reaction buffer and chilled on ice until addition. The remaining wells were treated with vehicle controls and fluorescence was recorded in the same manner. Readings from each set of three wells were averaged, and vehicle controls were subtracted from drug-treated wells. The resulting curves were smoothed by averaging four neighbors and smoothing to a second order polynomial to arrive at the displayed data. Average changes in fluorescence were calculated using data that were subjected to background subtraction, but not smoothed. In these cases, the reported values represent the absolute value of the difference between maximum (or, in the case of oligomycin, minimum) fluorescence after drug treatment and fluorescence just before drug treatment.

### Extraction of porphyrins and polar metabolites for LC-MS analysis

Parasites were harvested from HFFs when approximately half the vacuoles were lysed and prepared as for transcriptomics. Parasite pellets were quenched in 300 µl ice-cold 75% ACN supplemented with 1 µM deuteroporphyrin IX (Frontier Scientific, D510-9) and 0.5 µM isotopically labeled amino acids (Cambridge Isotopes, MSK-A2-1.2) and kept at -20 °C. Porphyrins were measured as previously described^99^ with the following modifications. To extract free porphyrins as well as polar metabolites, samples were sonicated for 10 cycles in a Bioruptor (Diagenode) at 4 °C with 30 sec on and 30 sec off, then incubated at 4 °C for 10 min. After a 10 min, 10.000 rpm spin in a table-top centrifuge (Eppendorf), the supernatant was collected and the pellet was washed with 100 µl 75% ACN and spun again. The supernatants from the two spins were combined. To extract protein-bound porphyrins, 300µl of 2:8 ratio of 1.7M HCl:ACN was added to the pellet, which was then vigorously shaken for 20 min at RT in a thermomixer (Eppendorf). After addition of 81 µl super-saturated MgSO_4_ solution and 24 µl 5 M NaCl, samples were vortexed for 30 sec and further shaken for 10 min at RT with occasional vortexing. Finally, after a 10 min 10,000 rpm spin in a tabletop centrifuge, the upper organic layer was collected. Extracted metabolites were dried in a CentriVap concentrator equipped with a cold trap (Labconco, Kansas City, MO) and reconstituted in an appropriate volume of 75% ACN (30–100 µL).

### LC-MS targeted analysis for polar metabolites

Polar samples were treated for small molecule LC-MS as previously described^100^. MS data acquisition on a QExactive benchtop orbitrap mass spectrometer equipped with an Ion Max source and a HESI II probe, which was coupled to a Dionex UltiMate 3000 UPLC system (Thermo Fisher Scientific, San Jose, CA) and was performed in a range of *m/z*= 70–1000, with the resolution set at 70,000, the AGC target at 1×10^6^, and the maximum injection time (Max IT) at 20 msec. For tSIM scans, the resolution was set at 70,000, the AGC target was 1×10^5^, and the max IT was 250 msec. Relative quantitation of polar metabolites was performed with XCalibur QuanBrowser 2.2 (Thermo Fisher Scientific) and TraceFinder software (ThermoFisher Scientific, Waltham, MA) using a 5 ppm mass tolerance and referencing an in-house library of chemical standards. Pooled samples and fractional dilutions were prepared as quality controls and only those metabolites were taken for further analysis, for which the correlation between the dilution factor and the peak area read was >0.95 (high confidence metabolites). Normalization for relative parasite amounts was based on the total integrated peak area values of high-confidence metabolites within an experimental batch after normalizing to the averaged factor from all mean-centered areas of the isotopically labeled amino acids internal standards. It is of note that normalizing to protein content by Bradford analysis or the sum of all area reads from high-confidence metabolites lead to similar results. The data were further Pareto transformed for MetaboAnalyst-based statistical or pathway analysis^101^.

### LC-MS targeted analysis of porphyrins

Free or protein-bound porphyrins were extracted as described above. Porphyrins were separated on a 2.6 µm, 150 x 3 mm C18 column (Phenomenex, 00F-4462-Y0) equipped with a 3.0 mm safe-guard column (Phenomenex, AJ0-8775) with a 22 min linear gradient of A: 0.1% formic acid in water and B: 0.1% formic acid in ACN at 0.7 mL/min flow. MS data acquisition on a triple quadrupole TSQ Quantum Access Max (Thermo Fisher Scientific) was in SRM mode with the following ion transitions: deuteroporphyrin IX 511.2 -> 451.95–452.45, protoporphyrin IX 563.2 -> 503.98–504.48, hemin 616.1 -> 556.9–557.4, and coproprotoporphyrin III or I 655.2 -> 594.9–595.4. Relative quantitation of porphyrins was performed with XCalibur QuanBrowser 2.2 (Thermo Fisher Scientific) using a 5 ppm mass tolerance and experimentally determined porphyrin retention times. Pooled samples and fractional dilutions were prepared as quality controls and only those measurements were taken for further analysis, for which the correlation between the dilution factor and the peak area read was >0.9. Total integrated peak area values were used as readout of relative concentrations after normalizing all values to the internal standard Deuteroporphyrin IX as well as to the averaged factor from all mean centered high confidence polar metabolites (see above polar metabolites analysis).

### *P. falciparum* maintenance and strain construction

*P. falciparum* asexual blood-stage parasites were cultured in human erythrocytes (3% hematocrit) and RPMI-1640 medium supplemented with 2 mM L-glutamine, 50 mg/L hypoxanthine, 25 mM HEPES, 0.225% NaHCO_3_, 10 mg/L gentamycin, and 0.5% (w/v) Albumax II (Invitrogen). Parasites were maintained at 37 °C in 5% O_2_, 5% CO_2_, and 90% N_2_. Cultures were stained with Giemsa, monitored by blood smears fixed in methanol, and viewed by light microscopy.

Transfections were performed by electroporating ring-stage parasites at 5–10% parasitemia with 50 μg of purified circular plasmid DNA resuspended in Cytomix^102^. One day after electroporation, parasites were exposed to 2.5 nM WR99210 for 6 days to select for transformed parasites. Parasite cultures were monitored by microscopy for up six weeks post electroporation, and recrudescent parasites were screened for editing by PCR. Positively-edited bulk cultures were cloned by limiting dilution in 96-well plates, and flow cytometry was used to screen for positive parasites after 17–20 days. Parasites were stained with 1 x SYBR Green (ThermoFisher) and 100 nM MitoTracker Deep Red (Invitrogen) and detected using an Accuri C6 flow cytometer (Becton Dickinson)^103^.

To generate the knockout construct for the serine protease *Pf* DegP (PF3D7_0807700) a donor sequence (834 bp total) was synthesized (Genewiz) fusing homology regions located in exon 2 (HR1) and exon 5 (HR2) with the addition of one nucleotide to introduce a frameshift in exon 5. The donor fragment was cloned into the pDC2 plasmid^104^ by amplifying with primers p15 and p16 (**Supplementary Table 9** using the restriction sites *EcoRI* and *AatII*. The pDC2 plasmid^105^ contains a codon-optimized Cas9 sequence under the regulatory control of the 5’ *calmodulin* (PF3D7_1434200) promoter and the 3’ *hsp86* (PF3D7_0708500) terminator as well as the *hdhfr* selectable marker which mediates resistance to the antiplasmodial agent WR99210 (Jacobus Pharmaceuticals). Two gRNA sequences (**Supplementary Table 9**) located in exon 4 were selected using chopchop^106^ and cloned into pDC2 using In-Fusion cloning kit (Takara Bio USA, Inc.). The final constructs pDC2-coCas9-*PfDegPKO*-*hdhfr* were transfected into Cam3.II^WT^ parasites as described above. Primers for diagnostic PCRs as well as sequencing primers are shown in **Figure 6b** and listed in **Supplementary Table 9**.

### Schizont-stage and ring-stage survival assays (SSA_38h_ and RSA_0–3h_)

These assays were carried out as previously described^37^, with minor modifications. In brief, parasite cultures were synchronized 1–2 times using 5% sorbitol (Fisher). For survival assays with succinyl acetone (SA; Sigma Aldrich), cultures were pretreated with 200 μM SA for 12 h prior to the onset of the ring-stage survival assay. Drug was kept on throughout the assay until assessment of parasite growth. Synchronous schizonts were incubated in RPMI-1640 containing 15 units/mL sodium heparin for 15 min at 37 °C to disrupt agglutinated erythrocytes, then concentrated over a gradient of 75% Percoll (Fisher) and washed once in RPMI-1640. For the schizont stage survival assay the concentrated schizonts were counted and seeded at 0.3% parasitemia followed by a 4 h exposure to 700 nM DHA or 0.1% DMSO (vehicle control). For the ring stage survival assay purified schizonts were incubated for 3 h with fresh RBCs to allow time for merozoite invasion. Cultures were subjected again to sorbitol treatment to eliminate remaining schizonts. 0–3 h post-invasion rings were adjusted to 1% parasitemia and 2% hematocrit and exposed to 700 nM DHA or 0.1% DMSO (vehicle control) for 4 h. Cells were washed to remove drug and returned to standard culture conditions for an additional 66 h. Parasite growth in each well was assessed using flow cytometry. Between 60,000 and 100,000 events were captured for each well. After 72 h, cultures generally expanded to 3–5 % parasitemia in DMSO-treated controls. Percent survival of DHA-treated parasites was calculated relative to the corresponding DMSO-treated control.

### Cellular heme fractionation assay

Heme fractionation was performed based on previously described methods^67,68^. Briefly, sorbitol-synchronized early ring-stage PfNF54 parasites (<5 h post-invasion) were incubated at 5% parasitemia and 2% hematocrit. Ring-stage parasites were harvested at 6 h and trophozoites were harvested at 20 h post incubation. RBCs were lysed with 0.05% saponin followed by multiple washes with PBS to remove traces of hemoglobin. The pellet was then resuspended in PBS and cell number quantified by flow cytometry. The remaining parasites were then lysed using hypotonic stress and sonication. Following centrifugation at 3600 rpm for 20 min at room temperature, treatment with 0.2 M HEPES buffer (pH 7.4), 4% SDS (w/v), 0.3 M NaCl, 25% pyridine, 0.3 M NaOH, 0.3 M HCl and distilled water, the fractions corresponding to hemoglobin, free heme and hemozoin were recovered as solution. The UV-visible spectrum of each heme fraction as an Fe(III)heme-pyridine complex was measured using a multi-well plate reader (Spectramax 340PC; Molecular Devices). The total amount of each heme species was quantified using a standard curve prepared from a standard solution of porcine hematin (Sigma-Aldrich) serially diluted in the same solvents used to process the pellets. The mass of each heme species in femtogram per cell (fg/cell) was then calculated by dividing the total amount of each heme species by the corresponding number of parasites. Statistical comparisons were made using one-way ANOVA (GraphPad Software Inc., La Jolla, CA, USA).

## ACRONYMS

α-KG: α-ketoglutarate
ALA: δ-aminolevulinic acid
ALAD: δ-aminolevulinic acid dehydrogenase
ALAS: δ-aminolevulinic acid synthase
ATQ: atovaquone
CPIII: coproporphyrinogen III
D3: Dulbecco’s modified Eagle’s medium containing 3% heat-inactivated fetal bovine serum
FAD: flavin adenine dinucleotide
Gly: glycine
HMB: hydroxymethylbilane
MS: Mass spectrometry
NADPH: nicotinamide adenine dinucleotide phosphate
NaFAc: sodium fluoroacetate
PGB: Porphobilinogen
PPO: protoporphyrinogen oxidase
PPIX: protoporphyrin IX
PPGIX: protoporphyrinogen IX
SA: succinyl acetone
SD: standard deviation
SEM: standard error of the mean
TCA: tricarboxylic acid cycle
TTFA: thenoyltrifluoroacetone

## SUPPLEMENTARY FIGURES

**Supplementary Figure 1.**
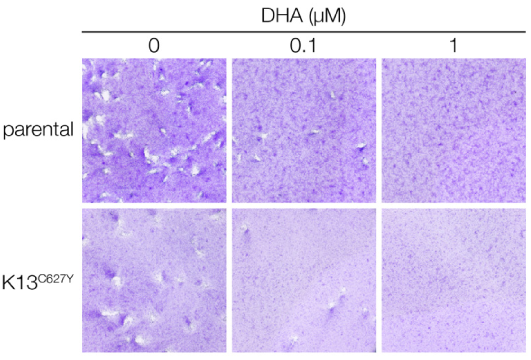
K13^C627Y^ parasites grow normally and do not show DHA resistance in plaque assay. Plaque assay or parental and K13^C627Y^ parasites, fixed after 7 days of drug treatment with the indicated concentration.

**Supplementary Figure 2.**
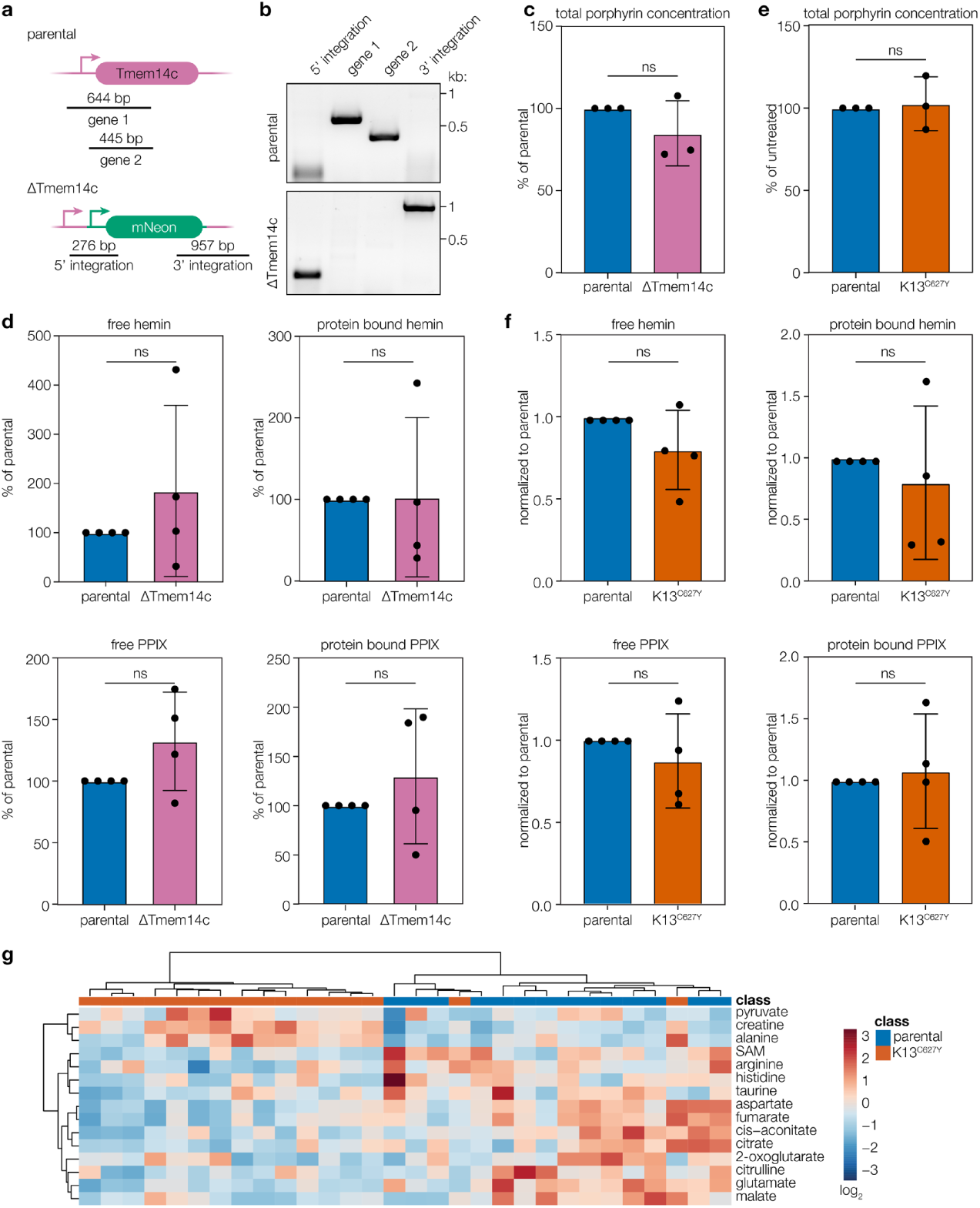
Construction of ΔTmem14c and porphyrin measurements from ΔTmem14c and K13^C627Y^. **a**, Schematic showing strategy for creating ΔTmem14c parasites. The coding sequence was replaced by an mNeonGreen cassette. **b**, PCR demonstrating correct deletion of *Tmem14c*. **c**, Total porphyrins were quantified from parental and ΔTmem14c parasites. Results are mean ± SD for n = 3 independent experiments, each performed in technical duplicate. **d**, Free or protein-bound hemin or PPIX were extracted from parental and ΔTmem14c parasites and quantified by LC-MS. Results are mean ± SD from n = 3 independent experiments, each performed in technical triplicate and normalized to untreated controls; *p* values are from two-way ANOVA; ns, not significant. **e**, Total porphyrins were quantified from parental and K13^C627Y^ parasites. Results are mean ± SD for n = independent experiments; ns, not significant by Student’s *t* test. **f** Free or protein-bound hemin or PPIX were extracted from parental and K13^C627Y^ parasites and quantified by LC-MS. Results are mean ± SD from n = 4 independent experiments, each performed at minimum in technical triplicate; ns, not significant by two-way ANOVA. **g**, MetaboAnalyst heatmap analysis of top 15 changed polar metabolites between parental and K13^C627Y^ parasites from 4 independent experiments. Metabolite peak areas were normalized to total metabolite signal (see **Methods**)

**Supplementary Figure 3.**
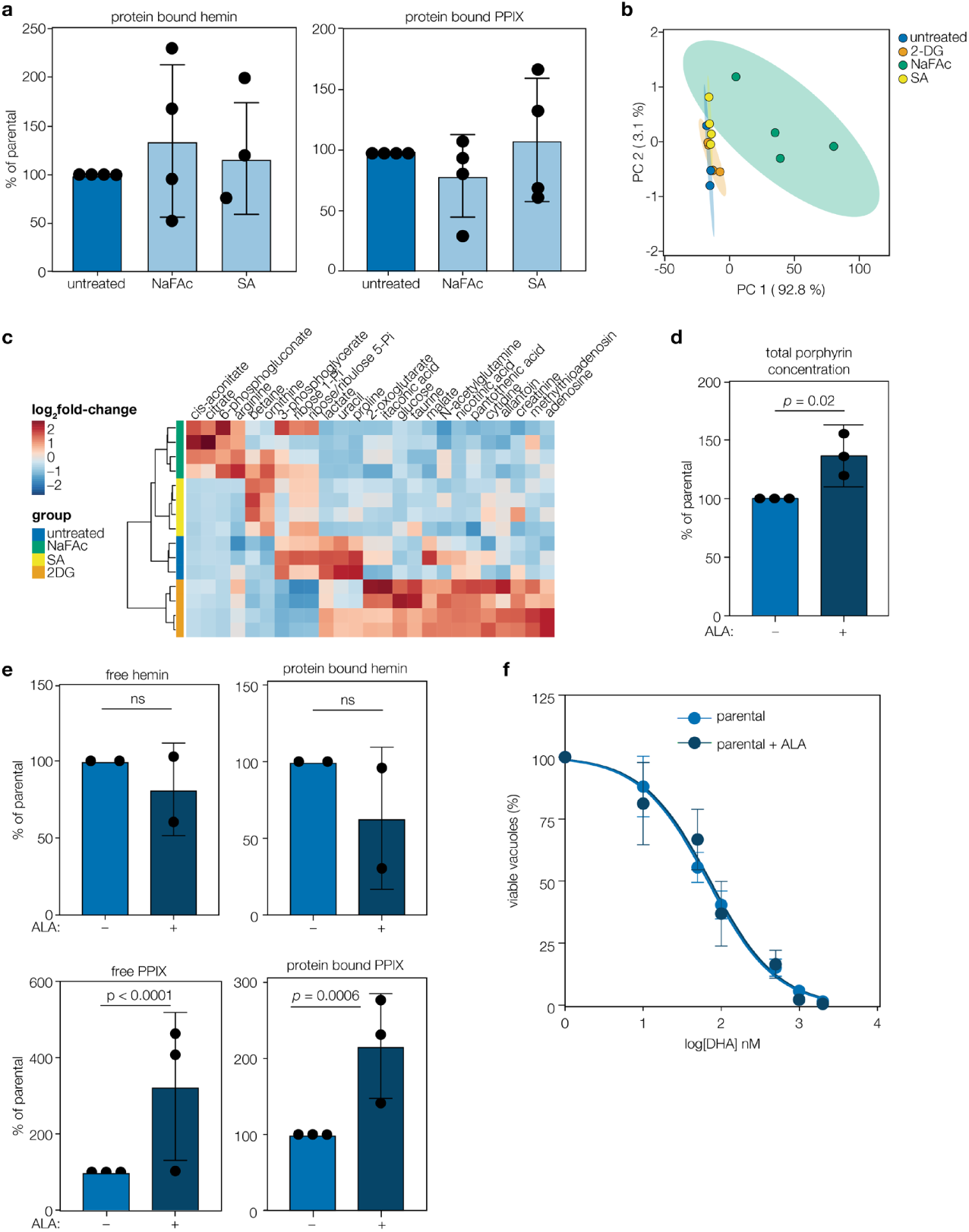
Effects of glycolysis, TCA, and heme biosynthesis modulators on parasite metabolism. **a**, Protein-bound hemin or protoporphyrin IX (PPIX) were extracted from parasites treated with the indicated compound, and quantified by LC-MS. Results are mean ± SD for n = 4 independent experiments, each performed at minimum in technical duplicate and normalised to the untreated control. ns, not significant by two-way ANOVA. **b**, PCA plot indicating variance of total polar metabolites between various compound-treated samples. 500 mM NaFAc treatment had significant global effects on metabolite abundance, while 5 mM 2-DG and 10 mM SA had more subtle effects and overlapped with the parental, untreated line. **c**, MetaboAnalyst heatmap analysis of the top 25 polar metabolites with altered abundance after the indicated treatment. Results are from a representative experiment performed in triplicate. Metabolite peak areas were normalized to total metabolite signal (see **Methods**). **d**, Normalized total porphyrin levels in untreated or 200 µM ALA treated parasites. Results are mean ± SD of n = 3 independent experiments performed in duplicate. *p* values from *t* test on raw values. **e**, Free or protein-bound hemin or PPIX were extracted from parasites treated with or without ALA and quantified by LC-MS. Results from three independent experiments (or 2 for heme measurements), normalized to untreated, performed in at least 3 replicates, ± SD. *p* values by two-way ANOVA; ns, not significant. **f**, Dose response curve of parental parasites or treated with ALA as above. No difference in the response to DHA could be seen (extra-sum-of-squares F test, *p* = 0.79). Results mean of at least three biological replicates, ± SEM.

**Supplementary Figure 4.**
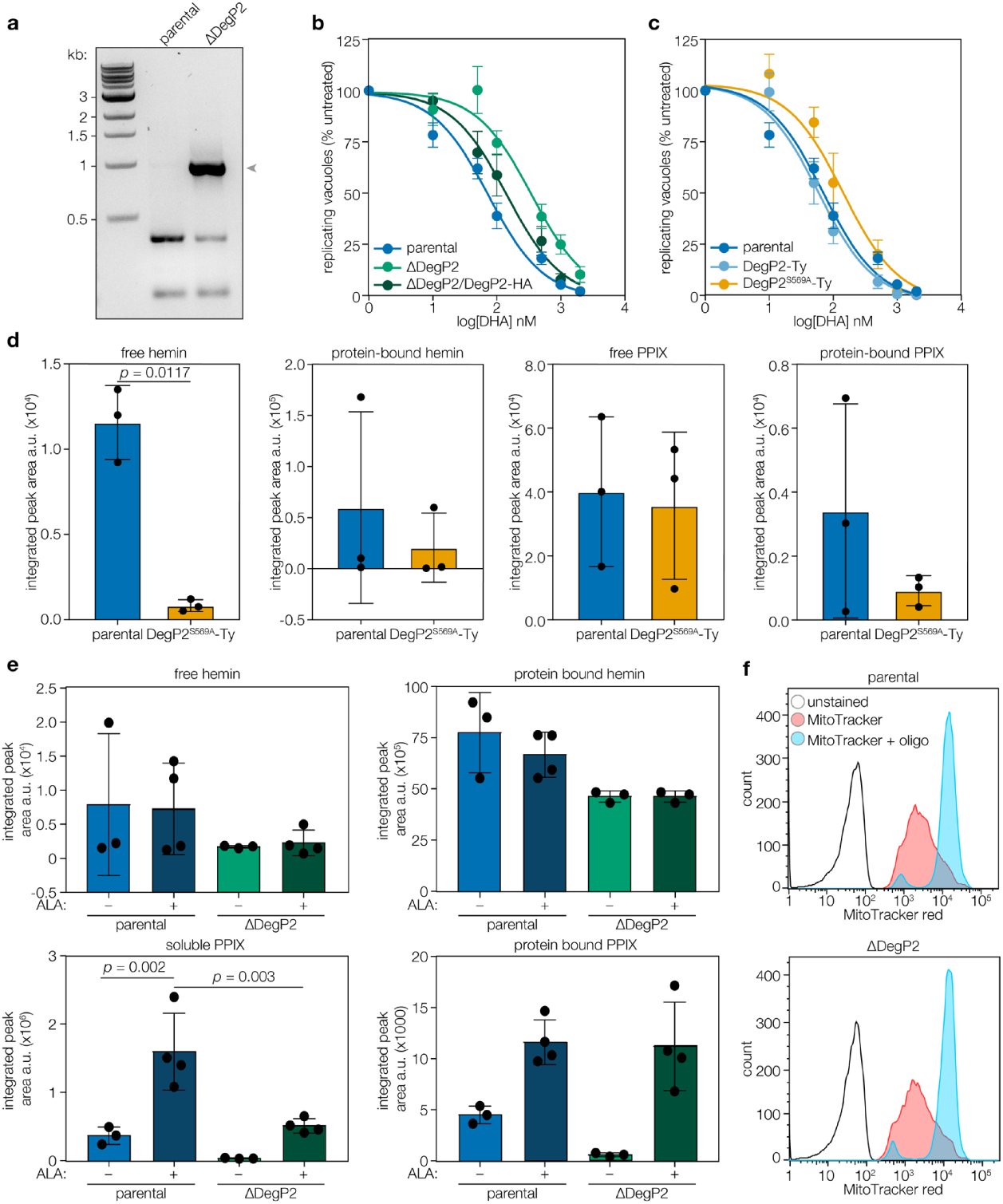
DegP2 has a role in DHA sensitivity and porphyrin biogenesis, which is dependent on its protease activity. **a**, PCR confirming loss of endogenous locus and replacement with YFP. Arrow indicates the size of the PCR product that’s expected after replacement. **b**, DHA dose-response curves. Results are mean ± SEM for n = 7, 7, or 4 independent experiments for the parental, ΔDegP2, or ΔDegP2/DegP2-HA strains, respectively. **c**, DHA dose-response curves. Results are mean ± SEM for n = 7, 4, or 4 independent replicates for the parental, DegP2-Ty, or DegP2^S569A^-Ty strains, respectively. **d**, Free or protein-bound hemin or PPIX were extracted from parental and DegP2^S569A^-Ty parasites and quantified by LC-MS. Results from one experiment performed in triplicate, ± SD. Significance tests by Student’s *t* test, no bar indicates not significant. **e**, Free or protein-bound hemin or PPIX were extracted from parental and ΔDegP2 parasites treated with or without ALA treatment and quantified by LC-MS. Results from one experiment performed in quadruplicate, ± SD. Significance tests by Student’s *t* test, no bar indicates not significant. **f**, Histograms of flow cytometry data of MitoTracker-stained parental and ΔDegP2 parasites, then treated or untreated with oligomycin. No differences were observed between parental and ΔDegP2 parasites. Results representative of two independent experiments.

## SUPPLEMENTARY TABLES & LEGENDS

**Supplementary Table 1. Summary of IC50 values.**

**Supplementary Table 2.** Full CRISPR screen data.

| Gene ID | Annotation | Drug score |  |  | Phenotype score<sup>a</sup> |
| --- | --- | --- | --- | --- | --- |
|  |  | Rep1 | Rep2 | Rep3 |  |
| TGGT1_290840 | serine protease (DegP2) | 9.97 | 3.90 | 7.03 | -1.87 |
| TGGT1_244200 | α-ketoglutarate dehydrogenase (E1) | 13.15 | 10.97 | 5.03 | -4.8 |
| TGGT1_219550 | α-ketoglutarate dehydrogenase (E2) | 12.01 | 12.26 | 4.99 | -4.25 |
| TGGT1_206470 | pyruvate dehydrogenase (PDH-E3II) | 12.90 | 12.55 | 9.07 | -3.23 |
| TGGT1_251680 | Translationally controlled tumor protein (TCTP) | ns | 2.65 | 3.24 | -1.37 |
| TGGT1_297080 | pyridoxal kinase | 5.63 | ns | 5.01 | -0.41 |
| TGGT1_272490 | protoporphyrinogen oxidase | 10.26 | ns | 9.66 | -3.87 |
| TGGT1_271410 | hypothetical protein | 2.59 | ns | 0.24 | -1.1 |

**Supplementary Table 3. Summary of top hits from three replicates of the genome-wide screen.** ns, not significantly enriched. ^a^(Sidik, 2016)

**Supplementary Table 4. Hits from MAGeCK analysis.** All genes significantly (defined as FDR <0.1) overrepresented using MAGeCK^43^ in each of the three independent replicates. Genes which appeared in more than one screen are highlighted.

**Supplementary Table 5. Raw data from untargeted polar metabolomics from parental lines treated with metabolic inhibitors (NaFAc, SA, 2-DG)**

**Supplementary Table 6.** MetaboAnalyst pathway analysis of polar metabolites from parental vs K13^C627Y^ parasites, normalized to mean parental values. Significantly different (FDR < 0.05) pathways are listed. Pathway impact values closer to 1 indicate higher node importance.

**Supplementary Table 7. RNAseq raw counts of parental and ΔDegP2 parasite lines.**

**Supplementary Table 8. Raw data from untargeted polar metabolomics from parental and ΔDegP2 parasite lines.**

**Supplementary Table 9. *T. gondii* and *P. falciparum* primers used in this study**

## REFERENCES

1. Tilley, L., Straimer, J., Gnädig, N. F., Ralph, S. A. & Fidock, D. A. Artemisinin Action and Resistance in Plasmodium falciparum. Trends Parasitol. 32, 682–696 (2016).

2. Konstat-Korzenny, E., Ascencio-Aragón, J. A., Niezen-Lugo, S. & Vázquez-López, R. Artemisinin and Its Synthetic Derivatives as a Possible Therapy for Cancer. Med Sci (Basel) 6, (2018).

3. Meshnick, S. R., Thomas, A., Ranz, A., Xu, C. M. & Pan, H. Z. Artemisinin (qinghaosu): the role of intracellular hemin in its mechanism of antimalarial action. Mol. Biochem. Parasitol. 49, 181–189 (1991).

4. Wang, J. et al. Haem-activated promiscuous targeting of artemisinin in Plasmodium falciparum. Nat. Commun. 6, 10111 (2015).

5. Ismail, H. M. et al. Artemisinin activity-based probes identify multiple molecular targets within the asexual stage of the malaria parasites Plasmodium falciparum 3D7. Proc. Natl. Acad. Sci. U. S. A. 113, 2080–2085 (2016).

6. Hartwig, C. L. et al. Accumulation of artemisinin trioxane derivatives within neutral lipids of Plasmodium falciparum malaria parasites is endoperoxide-dependent. Biochem. Pharmacol. 77, 322–336 (2009).

7. Heller, L. E., Goggins, E. & Roepe, P. D. Dihydroartemisinin–Ferriprotoporphyrin IX Adduct Abundance in Plasmodium falciparum Malarial Parasites and the Relationship to Emerging Artemisinin Resistance. Biochemistry 57, 6935–6945 (2018).

8. Dondorp, A. M. et al. Artemisinin resistance in Plasmodium falciparum malaria. N. Engl. J. Med. 361, 455–467 (2009).

9. Noedl, H. et al. Evidence of artemisinin-resistant malaria in western Cambodia. N. Engl. J. Med. 359, 2619–2620 (2008).

10. van der Pluijm, R. W. et al. Determinants of dihydroartemisinin-piperaquine treatment failure in Plasmodium falciparum malaria in Cambodia, Thailand, and Vietnam: a prospective clinical, pharmacological, and genetic study. Lancet Infect. Dis. (2019).

11. White, N. J. et al. Malaria. Lancet 383, 723–735 (2014).

12. Ariey, F. et al. A molecular marker of artemisinin-resistant Plasmodium falciparum malaria. Nature 505, 50–55 (2014).

13. Straimer, J. et al. K13-propeller mutations confer artemisinin resistance in Plasmodium falciparum clinical isolates. Science 347, 428–431 (2015).

14. Dogovski, C. et al. Targeting the cell stress response of Plasmodium falciparum to overcome artemisinin resistance. PLoS Biol. 13, e1002132 (2015).

15. Mok, S. et al. Drug resistance. Population transcriptomics of human malaria parasites reveals the mechanism of artemisinin resistance. Science 347, 431–435 (2015).

16. Breglio, K. F. et al. A single nucleotide polymorphism in the Plasmodium falciparum atg18 gene associates with artemisinin resistance and confers enhanced parasite survival under nutrient deprivation. Malar. J. 17, 391 (2018).

17. Mbengue, A. et al. A molecular mechanism of artemisinin resistance in Plasmodium falciparum malaria. Nature 520, 683–687 (2015).

18. Demas, A. R. et al. Mutations in Plasmodium falciparum actin-binding protein coronin confer reduced artemisinin susceptibility. Proc. Natl. Acad. Sci. U. S. A. 115, 12799–12804 (2018).

19. Henriques, G. et al. Artemisinin resistance in rodent malaria--mutation in the AP2 adaptor μ-chain suggests involvement of endocytosis and membrane protein trafficking. Malar. J. 12, 118 (2013).

20. Rocamora, F. et al. Oxidative stress and protein damage responses mediate artemisinin resistance in malaria parasites. PLoS Pathog. 14, e1006930 (2018).

21. Henrici, R. C. et al. Modification of an atypical clathrin-independent AP-2 adaptin complex of Plasmodium falciparum reduces susceptibility to artemisinin. bioRxiv (2019). doi:10.1101/621078

22. Xie, S. C. et al. Haemoglobin degradation underpins the sensitivity of early ring stage Plasmodium falciparum to artemisinins. J. Cell Sci. 129, 406–416 (2016).

23. Mukherjee, A. et al. Artemisinin resistance without pfkelch13 mutations in Plasmodium falciparum isolates from Cambodia. Malar. J. 16, 195 (2017).

24. Sutherland, C. J. et al. pfk13-Independent Treatment Failure in Four Imported Cases of Plasmodium falciparum Malaria Treated with Artemether-Lumefantrine in the United Kingdom. Antimicrob. Agents Chemother. 61, (2017).

25. 26. Berens, R. L., Krug, E. C., Nash, P. B. & Curiel, T. J. Selection and characterization of Toxoplasma gondii mutants resistant to artemisinin. J. Infect. Dis. 177, 1128–1131 (1998).

26. Reynolds, M. G., Oh, J. & Roos, D. S. In vitro generation of novel pyrimethamine resistance mutations in the Toxoplasma gondii dihydrofolate reductase. Antimicrob. Agents Chemother. 45, 1271–1277 (2001).

27. Cowell, A. N. et al. Mapping the malaria parasite druggable genome by using in vitro evolution and chemogenomics. Science 359, 191–199 (2018).

28. Shalem, O. et al. Genome-scale CRISPR-Cas9 knockout screening in human cells. Science 343, 84–87 (2014).

29. Hou, P. et al. A Genome-Wide CRISPR Screen Identifies Genes Critical for Resistance to FLT3 Inhibitor AC220. Cancer Res. 77, 4402–4413 (2017).

30. Shi, C.-X. et al. CRISPR Genome-Wide Screening Identifies Dependence on the Proteasome Subunit PSMC6 for Bortezomib Sensitivity in Multiple Myeloma. Mol. Cancer Ther. 16, 2862–2870 (2017).

31. Sidik, S. M., Huet, D. & Lourido, S. CRISPR-Cas9-based genome-wide screening of Toxoplasma gondii. Nature Protocols 13, 307–323 (2018).

32. Sidik, S. M. et al. A Genome-wide CRISPR Screen in Toxoplasma Identifies Essential Apicomplexan Genes. Cell 166, 1423–1435.e12 (2016).

33. Touquet, B. et al. High-content imaging assay to evaluate Toxoplasma gondii infection and proliferation: A multiparametric assay to screen new compounds. PLoS One 13, e0201678 (2018).

34. Parapini, S., Olliaro, P., Navaratnam, V., Taramelli, D. & Basilico, N. Stability of the antimalarial drug dihydroartemisinin under physiologically relevant conditions: implications for clinical treatment and pharmacokinetic and in vitro assays. Antimicrob. Agents Chemother. 59, 4046–4052 (2015).

35. Ghorbal, M. et al. Genome editing in the human malaria parasite Plasmodium falciparum using the CRISPR-Cas9 system. Nat. Biotechnol. 32, 819–821 (2014).

36. Wang, Z. et al. Artemisinin resistance at the China-Myanmar border and association with mutations in the K13 propeller gene. Antimicrob. Agents Chemother. 59, 6952–6959 (2015).

37. Witkowski, B. et al. Novel phenotypic assays for the detection of artemisinin-resistant Plasmodium falciparum malaria in Cambodia: in-vitro and ex-vivo drug-response studies. Lancet Infect. Dis. 13, 1043–1049 (2013).

38. Klonis, N. et al. Altered temporal response of malaria parasites determines differential sensitivity to artemisinin. Proc. Natl. Acad. Sci. U. S. A. 110, 5157–5162 (2013).

39. Radke, J. R. et al. Defining the cell cycle for the tachyzoite stage of Toxoplasma gondii. Mol. Biochem. Parasitol. 115, 165–175 (2001).

40. Markus, B. M., Bell, G. W., Lorenzi, H. A. & Lourido, S. Optimizing Systems for Cas9 Expression in Toxoplasma gondii. mSphere 4, (2019).

41. Zimmermann, L. et al. A Completely Reimplemented MPI Bioinformatics Toolkit with a New HHpred Server at its Core. J. Mol. Biol. 430, 2237–2243 (2018).

42. Yien, Y. Y. et al. TMEM14C is required for erythroid mitochondrial heme metabolism. J. Clin. Invest. 124, 4294–4304 (2014).

43. Li, W. et al. MAGeCK enables robust identification of essential genes from genome-scale CRISPR/Cas9 knockout screens. Genome Biol. 15, 554 (2014).

44. Astner, I. et al. Crystal structure of 5-aminolevulinate synthase, the first enzyme of heme biosynthesis, and its link to XLSA in humans. EMBO J. 24, 3166–3177 (2005).

45. MacRae, J. I. et al. Mitochondrial metabolism of glucose and glutamine is required for intracellular growth of Toxoplasma gondii. Cell Host Microbe 12, 682–692 (2012).

46. Shanmugam, D., Wu, B., Ramirez, U., Jaffe, E. K. & Roos, D. S. Plastid-associated Porphobilinogen Synthase fromToxoplasma gondii. Journal of Biological Chemistry 285, 22122–22131 (2010).

47. Huet, D., Rajendran, E., van Dooren, G. G. & Lourido, S. Identification of cryptic subunits from an apicomplexan ATP synthase. Elife 7, (2018).

48. Blume, M. et al. Host-derived glucose and its transporter in the obligate intracellular pathogen Toxoplasma gondii are dispensable by glutaminolysis. Proc. Natl. Acad. Sci. U. S. A. 106, 12998–13003 (2009).

49. Sinclair, P. R., Gorman, N. & Jacobs, J. M. Measurement of Heme Concentration. Curr. Protoc. Toxicol. 00, 8.3.1–8.3.7 (1999).

50. Klonis, N. et al. Artemisinin activity against Plasmodium falciparum requires hemoglobin uptake and digestion. Proc. Natl. Acad. Sci. U. S. A. 108, 11405–11410 (2011).

51. Fujiwara, T. et al. Exploring the Potential Usefulness of 5-Aminolevulinic Acid for X-Linked Sideroblastic Anemia. Blood 124, 215–215 (2014).

52. Zhang, S. & Gerhard, G. S. Heme mediates cytotoxicity from artemisinin and serves as a general anti-proliferation target. PLoS One 4, e7472 (2009).

53. Sigala, P. A., Crowley, J. R., Henderson, J. P. & Goldberg, D. E. Deconvoluting heme biosynthesis to target blood-stage malaria parasites. Elife 4, (2015).

54. Lentini, G. et al. Characterization of Toxoplasma DegP, a rhoptry serine protease crucial for lethal infection in mice. PLoS One 12, e0189556 (2017).

55. Carter, P. & Wells, J. A. Dissecting the catalytic triad of a serine protease. Nature 332, 564–568 (1988).

56. Hedstrom, L. Serine protease mechanism and specificity. Chem. Rev. 102, 4501–4524 (2002).

57. Sun, R. et al. Crystal structure of Arabidopsis Deg2 protein reveals an internal PDZ ligand locking the hexameric resting state. J. Biol. Chem. 287, 37564–37569 (2012).

58. Gardner, M. J. et al. Mitochondrial DNA of the human malarial parasite Plasmodium falciparum. Mol. Biochem. Parasitol. 31, 11–17 (1988).

59. Trawick, J. D., Wright, R. M. & Poyton, R. O. Transcription of yeast COX6, the gene for cytochrome c oxidase subunit VI, is dependent on heme and on the HAP2 gene. J. Biol. Chem. 264, 7005–7008 (1989).

60. Vijayasarathy, C., Damle, S., Lenka, N. & Avadhani, N. G. Tissue variant effects of heme inhibitors on the mouse cytochrome c oxidase gene expression and catalytic activity of the enzyme complex. Eur. J. Biochem. 266, 191–200 (1999).

61. Sun, F. et al. Crystal structure of mitochondrial respiratory membrane protein complex II. Cell 121, 1043–1057 (2005).

62. Biagini, G. A., Viriyavejakul, P., O’neill, P. M., Bray, P. G. & Ward, S. A. Functional characterization and target validation of alternative complex I of Plasmodium falciparum mitochondria. Antimicrob. Agents Chemother. 50, 1841–1851 (2006).

63. Syafruddin, D., Siregar, J. E. & Marzuki, S. Mutations in the cytochrome b gene of Plasmodium berghei conferring resistance to atovaquone. Mol. Biochem. Parasitol. 104, 185–194 (1999).

64. Korsinczky, M. et al. Mutations in Plasmodium falciparumCytochrome b That Are Associated with Atovaquone Resistance Are Located at a Putative Drug-Binding Site. Antimicrob. Agents Chemother. 44, 2100–2108 (2000).

65. Vercesi, A. E., Rodrigues, C. O., Uyemura, S. A., Zhong, L. & Moreno, S. N. Respiration and oxidative phosphorylation in the apicomplexan parasite Toxoplasma gondii. J. Biol. Chem. 273, 31040–31047 (1998).

66. Krumschnabel, G., Eigentler, A., Fasching, M. & Gnaiger, E. Chapter Nine - Use of Safranin for the Assessment of Mitochondrial Membrane Potential by High-Resolution Respirometry and Fluorometry. in Methods in Enzymology (eds. Galluzzi, L. & Kroemer, G.) 542, 163–181 (Academic Press, 2014).

67. Combrinck, J. M. et al. Optimization of a multi-well colorimetric assay to determine haem species in Plasmodium falciparum in the presence of anti-malarials. Malar. J. 14, 253 (2015).

68. Combrinck, J. M. et al. Insights into the role of heme in the mechanism of action of antimalarials. ACS Chem. Biol. 8, 133–137 (2013).

69. Zhang, S. & Gerhard, G. S. Heme activates artemisinin more efficiently than hemin, inorganic iron, or hemoglobin. Bioorg. Med. Chem. 16, 7853–7861 (2008).

70. Miksanova, M. et al. Characterization of heme-regulated eIF2alpha kinase: roles of the N-terminal domain in the oligomeric state, heme binding, catalysis, and inhibition. Biochemistry 45, 9894–9905 (2006).

71. Hanna, D. A. et al. Heme dynamics and trafficking factors revealed by genetically encoded fluorescent heme sensors. Proc. Natl. Acad. Sci. U. S. A. 113, 7539–7544 (2016).

72. 73. van Dooren, G. G., Kennedy, A. T. & McFadden, G. I. The use and abuse of heme in apicomplexan parasites. Antioxid. Redox Signal. 17, 634–656 (2012).

73. Shanmugasundram, A., Gonzalez-Galarza, F. F., Wastling, J. M., Vasieva, O. & Jones, A. R. Library of Apicomplexan Metabolic Pathways: a manually curated database for metabolic pathways of apicomplexan parasites. Nucleic Acids Res. 41, D706–13 (2013).

74. Giacometti, A., Cirioni, O. & Scalise, G. In-vitro activity of macrolides alone and in combination with artemisin, atovaquone, dapsone, minocycline or pyrimethamine against Cryptosporidium parvum. J. Antimicrob. Chemother. 38, 399–408 (1996).

75. Heller, L. E. & Roepe, P. D. Quantification of Free Ferriprotoporphyrin IX Heme and Hemozoin for Artemisinin Sensitive versus Delayed Clearance Phenotype Plasmodium falciparum Malarial Parasites. Biochemistry 57, 6927–6934 (2018).

76. Skinner, T. S., Manning, L. S., Johnston, W. A. & Davis, T. M. In vitro stage-specific sensitivity of Plasmodium falciparum to quinine and artemisinin drugs. Int. J. Parasitol. 26, 519–525 (1996).

77. Delves, M. et al. The activities of current antimalarial drugs on the life cycle stages of Plasmodium: a comparative study with human and rodent parasites. PLoS Med. 9, e1001169 (2012).

78. Pomel, S., Luk, F. C. Y. & Beckers, C. J. M. Host cell egress and invasion induce marked relocations of glycolytic enzymes in Toxoplasma gondii tachyzoites. PLoS Pathog. 4, e1000188 (2008).

79. Abshire, J. R., Rowlands, C. J., Ganesan, S. M., So, P. T. C. & Niles, J. C. Quantification of labile heme in live malaria parasites using a genetically encoded biosensor. Proc. Natl. Acad. Sci. U. S. A. 114, E2068–E2076 (2017).

80. Abu Bakar, N., Klonis, N., Hanssen, E., Chan, C. & Tilley, L. Digestive-vacuole genesis and endocytic processes in the early intraerythrocytic stages of Plasmodium falciparum. J. Cell Sci. 123, 441–450 (2010).

81. Schuhmann, H. & Adamska, I. Deg proteases and their role in protein quality control and processing in different subcellular compartments of the plant cell. Physiol. Plant. 145, 224–234 (2012).

82. Chang, Z. The function of the DegP (HtrA) protein: Protease versus chaperone. IUBMB Life 68, 904–907 (2016).

83. Gao, T. & O’Brian, M. R. Control of DegP-dependent degradation of c-type cytochromes by heme and the cytochrome c maturation system in Escherichia coli. J. Bacteriol. 189, 6253–6259 (2007).

84. Sharma, S., Jadli, M., Singh, A., Arora, K. & Malhotra, P. A secretory multifunctional serine protease, DegP of Plasmodium falciparum, plays an important role in thermo-oxidative stress, parasite growth and development. FEBS J. 281, 1679–1699 (2014).

85. Lenka, N., Vijayasarathy, C., Mullick, J. & Avadhani, N. G. Structural organization and transcription regulation of nuclear genes encoding the mammalian cytochrome c oxidase complex. Prog. Nucleic Acid Res. Mol. Biol. 61, 309–344 (1998).

86. Fontanesi, F., Soto, I. C., Horn, D. & Barrientos, A. Assembly of mitochondrial cytochrome c-oxidase, a complicated and highly regulated cellular process. Am. J. Physiol. Cell Physiol. 291, C1129–47 (2006).

87. Soto, I. C., Fontanesi, F., Myers, R. S., Hamel, P. & Barrientos, A. A heme-sensing mechanism in the translational regulation of mitochondrial cytochrome c oxidase biogenesis. Cell Metab. 16, 801–813 (2012).

88. Huynh, M.-H. & Carruthers, V. B. Tagging of endogenous genes in a Toxoplasma gondii strain lacking Ku80. Eukaryot. Cell 8, 530–539 (2009).

89. Sidik, S. M., Hackett, C. G., Tran, F., Westwood, N. J. & Lourido, S. Efficient genome engineering of Toxoplasma gondii using CRISPR/Cas9. PLoS One 9, e100450 (2014).

90. Gibson, D. G. et al. Enzymatic assembly of DNA molecules up to several hundred kilobases. Nat. Methods 6, 343–345 (2009).

91. Bastin, P., Bagherzadeh, Z., Matthews, K. R. & Gull, K. A novel epitope tag system to study protein targeting and organelle biogenesis in Trypanosoma brucei. Mol. Biochem. Parasitol. 77, 235–239 (1996).

92. Lourido, S. et al. Calcium-dependent protein kinase 1 is an essential regulator of exocytosis in Toxoplasma. Nature 465, 359–362 (2010).

93. Sidik, S. M. et al. Using a Genetically Encoded Sensor to Identify Inhibitors of Toxoplasma gondii Ca2+ Signaling. J. Biol. Chem. 291, 9566–9580 (2016).

94. Gajria, B. et al. ToxoDB: an integrated Toxoplasma gondii database resource. Nucleic Acids Res. 36, D553–6 (2008).

95. Pino, P. et al. Dual targeting of antioxidant and metabolic enzymes to the mitochondrion and the apicoplast of Toxoplasma gondii. PLoS Pathog. 3, e115 (2007).

96. Schindelin, J. et al. Fiji: an open-source platform for biological-image analysis. Nat. Methods 9, 676–682 (2012).

97. Dobin, A. et al. STAR: ultrafast universal RNA-seq aligner. Bioinformatics 29, 15–21 (2013).

98. Love, M. I., Huber, W. & Anders, S. Moderated estimation of fold change and dispersion for RNA-seq data with DESeq2. Genome Biol. 15, 550 (2014).

99. Fyrestam, J. & Östman, C. Determination of heme in microorganisms using HPLC-MS/MS and cobalt(III) protoporphyrin IX inhibition of heme acquisition in Escherichia coli. Analytical and Bioanalytical Chemistry 409, 6999–7010 (2017).

100. Birsoy, K. et al. An Essential Role of the Mitochondrial Electron Transport Chain in Cell Proliferation Is to Enable Aspartate Synthesis. Cell 162, 540–551 (2015).

101. Xia, J., Sinelnikov, I. V., Han, B. & Wishart, D. S. MetaboAnalyst 3.0—making metabolomics more meaningful. Nucleic Acids Res. 43, W251–W257 (2015).

102. Fidock, D. A., Nomura, T. & Wellems, T. E. Cycloguanil and its parent compound proguanil demonstrate distinct activities against Plasmodium falciparum malaria parasites transformed with human dihydrofolate reductase. Mol. Pharmacol. 54, 1140–1147 (1998).

103. Ekland, E. H., Schneider, J. & Fidock, D. A. Identifying apicoplast-targeting antimalarials using high-throughput compatible approaches. FASEB J. 25, 3583–3593 (2011).

104. Fidock, D. A. et al. Mutations in the P. falciparum digestive vacuole transmembrane protein PfCRT and evidence for their role in chloroquine resistance. Mol. Cell 6, 861–871 (2000).

105. Lee, M. C. S., Moura, P. A., Miller, E. A. & Fidock, D. A. Plasmodium falciparum Sec24 marks transitional ER that exports a model cargo via a diacidic motif. Mol. Microbiol. 68, 1535–1546 (2008).

106. Labun, K. et al. CHOPCHOP v3: expanding the CRISPR web toolbox beyond genome editing. Nucleic Acids Res. 47, W171–W174 (2019).

